# NAD(H) homeostasis is essential for host protection mediated by glycolytic myeloid cells in tuberculosis

**DOI:** 10.1101/2022.09.11.507472

**Authors:** Hayden T. Pacl, Krishna C. Chinta, Vineel P. Reddy, Sajid Nadeem, Ritesh R. Sevalkar, Kierveshan Nargan, Kapongo Lumamba, Threnesan Naidoo, Joel N. Glasgow, Anupam Agarwal, Adrie J. C. Steyn

## Abstract

*Mycobacterium tuberculosis* (*Mtb*) disrupts glycolytic flux in infected myeloid cells through an unclear mechanism. Flux through the glycolytic pathway in myeloid cells is inextricably linked to the availability of NAD^+^, which is maintained by NAD^+^ salvage and lactate metabolism. Using lung tissue from tuberculosis (TB) patients and myeloid deficient LDHA (*Ldha*^*LysM−/−*^) mice, we demonstrate that glycolysis in myeloid cells is essential for protective immunity in TB. Glycolytic myeloid cells are essential for the early recruitment of multiple classes of immune cells and the protective effects of IFNγ. We identified NAD^+^ depletion as central to the glycolytic inhibition caused by *Mtb*. Lastly, we show that the NAD^+^ precursor nicotinamide exerts a host-directed, antimycobacterial effect, and that nicotinamide prophylaxis and treatment reduces *Mtb* lung burden *in vivo*. These findings provide new insight into how *Mtb* alters host metabolism through perturbation of NAD(H) homeostasis and reprogramming of glycolysis, highlighting this pathway as a potential therapeutic target.

## Introduction

*Mycobacterium tuberculosis* (*Mtb*), the bacterium that causes tuberculosis (TB), remains a leading cause of death worldwide despite curative antibiotic therapy^1^. The success of *Mtb* as a pathogen is attributable to an array of virulence factors that modulate the host immune response to allow escape from host phagocytes, delayed onset of adaptive immunity, and the chronic inflammation and immunopathology characteristic of TB^2^. Several virulence factors have been well characterized since *Mtb* was first isolated in 1882; however, our understanding of the immunomodulatory strategy of this pathogen remains incomplete. A growing body of evidence indicates that host immunometabolism, the intrinsic link between metabolism and immune function, is an important component of TB pathogenesis^3–5^. In this regard, the metabolism of myeloid cells has emerged as a crucial determinant of TB outcomes, given that myeloid cells are the primary reservoir for *Mtb* in the infected host^6^ and their inflammatory functions are directly responsible for controlling infection and the immunopathology that defines TB^7–10^. Within this subset, sulfur^11^ and iron^12,13^ metabolism as well as central carbon metabolism^14^ play a role in host protection in TB.

Glycolysis is of particular interest^14–20^ as it is the pathway in central carbon metabolism by which cells oxidize glucose to pyruvate, simultaneously converting ADP and NAD^+^ to ATP and NADH, respectively. While glycolysis is often thought of as coupled with mitochondrial respiration under aerobic conditions, myeloid cells maintain high levels of flux through this anaerobic pathway even in the presence of oxygen to drive inflammation across many systems, referred to as aerobic glycolysis or the Warburg effect^14,21^. Initial studies using nonvirulent *Mtb* (*i*.*e*., killed or attenuated) suggest that *Mtb* infection prompts macrophages, a myeloid subset, to carry out aerobic glycolysis^22,23^, but a growing body of literature indicates that live, virulent *Mtb* decreases the glycolytic capacity of macrophages following infection^16,19,20^. Other studies associating glycolytic flux in macrophages with bacillary control both *in vitro* and *in vivo* suggest the consequences of impaired glycolytic capacity in these cells^15,17,18^. However, causally implicating glycolysis in these studies has been confounded by using experimental manipulations that lead to large-scale disruption of glycolysis and connected pathways, such as the use of 2-deoxy-*D*-glucose (2DG), an inhibitor of glucose uptake, and deletion of *Hif1A*, the gene encoding the highly conserved transcription factor, hypoxia inducible factor-1α (HIF1α). 2DG is also a competitive inhibitor of hexokinase at the start of glycolysis. Consequences of this inhibition are dysregulation of the linked pentose phosphate pathway (PPP) and the TCA cycle, which uses the end-product of glycolysis, *i*.*e*., pyruvate, as a substrate. Similarly, HIF1α has been shown to broadly regulate glycolysis, and unsurprisingly, deletion of *Hif1A* triggers a large pleotropic effect which confounds interpretation of results^24^. Therefore, an experimental approach that selectively targets a distinct step in glycolysis during *Mtb* infection would provide compelling evidence for its role in the control of TB.

Glycolysis is an essential metabolic process and is regulated at multiple levels. One cell-intrinsic approach to regulating glycolytic flux would be to modulate the availability of NAD^+^, the electron acceptor in glycolysis. In inflammatory myeloid cells, NAD^+^ availability [*i*.*e*., NAD(H) homeostasis] is largely dependent on: (i) maintaining the overall abundance of NAD(H) via the NAD^+^ salvage pathway, and (ii) regenerating NAD^+^ from NADH via lactate fermentation. NAD^+^ salvage regenerates NAD^+^ from the precursor nicotinamide (NAM), which is also a product of NAD^+^-consuming enzymes that are increased in inflammatory myeloid cells^25^. Lactate metabolism, on the other hand, couples the oxidation of NADH to NAD^+^ with the reduction of pyruvate to lactate. The reversible process of lactate fermentation is catalyzed by lactate dehydrogenase (LDH), a tetramer composed of LDHA and LDHB subunits. When the tetramer is composed predominantly (or exclusively) of LDHA subunits, the subunit predominantly expressed in myeloid cells, it preferentially converts pyruvate to lactate, and NADH to NAD^+^. However, LDHB subunits favor the opposite reaction. Thus, the intrinsic link between NAD(H) homeostasis and glycolysis provides several targets for the highly selective manipulation of host-cell glycolytic flux, improving on the frequent use of 2DG that severely impairs glycolysis, oxidative phosphorylation (OXPHOS), the PPP, and the TCA cycle.

Given the unmet need for shorter, simpler, or more tolerable regimens for TB treatment, specific manipulation of host glycolysis represents a potential host-directed therapy (HDT). This approach would supplement existing TB antibiotic therapy by enhancing the microbicidal function and/or limiting inflammation caused by host immune cells, especially myeloid cells^26–28^. Thus, the explicit contribution of glycolytic capacity in myeloid cells to host protection against TB is unknown. We therefore tested the hypothesis that NAD(H)-mediated glycolytic flux in myeloid cells is essential for host protection in TB. We examined LDHA protein expression patterns in human TB lung tissue, employed a myeloid-specific *Ldha* knockout mouse (*Ldha*^*LysM*−/−^) that exhibits reduced glycolytic capacity in myeloid cells^29,30^, and performed a series of bioenergetic and metabolomic experiments in *Ldha*^*LysM*−/−^ macrophages. We applied these findings to *in vitro* and *in vivo* models of *Mtb* infection using a pharmacological inhibitor and nutritional supplementation. Our findings highlight the essential role for glycolytic myeloid cells in host protection and the mechanism by which *Mtb* disrupts glycolysis in these cells. Further, increasing the glycolytic capacity of myeloid cells in the context of TB could represent a new and innovative approach to the prevention and treatment of TB.

## Results

### LDHA expression within the spectrum of human tuberculosis

Given the complexity and diversity of lesions in pulmonary tuberculosis and the suboptimal representation of this spectrum within available animal models^31,32^, we examined LDHA expression patterns in lung tissue resected from patients with TB to determine its clinical relevance across the histopathological spectrum of human pulmonary TB. In non-necrotizing granulomas (NNGs), we observed clusters of LDHA-positive cells with a lower nucleus:cytoplasm ratio, likely macrophages, and some lymphocytes, spread throughout a predominantly lymphocytic infiltrate (yellow arrows; Figure 1A). In early necrotizing granulomas, we observed intense, homogenous LDHA staining of the necrotic core (Figure 1B), representing LDHA released from necrotic immune cells. Heterogenous staining for LDHA is also present throughout areas of granulomatous inflammation and the developing necrotic core of more advanced necrotic lesions (Figure 1C). Lastly, in established, necrotic granulomas, LDHA staining is limited to the granulation layer (Figure 1D) and proximal regions of granulomatous inflammation. This stands in contrast to early necrotic granulomas, which exhibit robust staining of necrotic debris and illustrate a pattern consistent with a loss of staining with increased age of the lesion. Thus, LDHA expression, and consequently an increased capacity for lactate fermentation, is localized to regions of active inflammation within the human TB lung.

**Figure 1.**
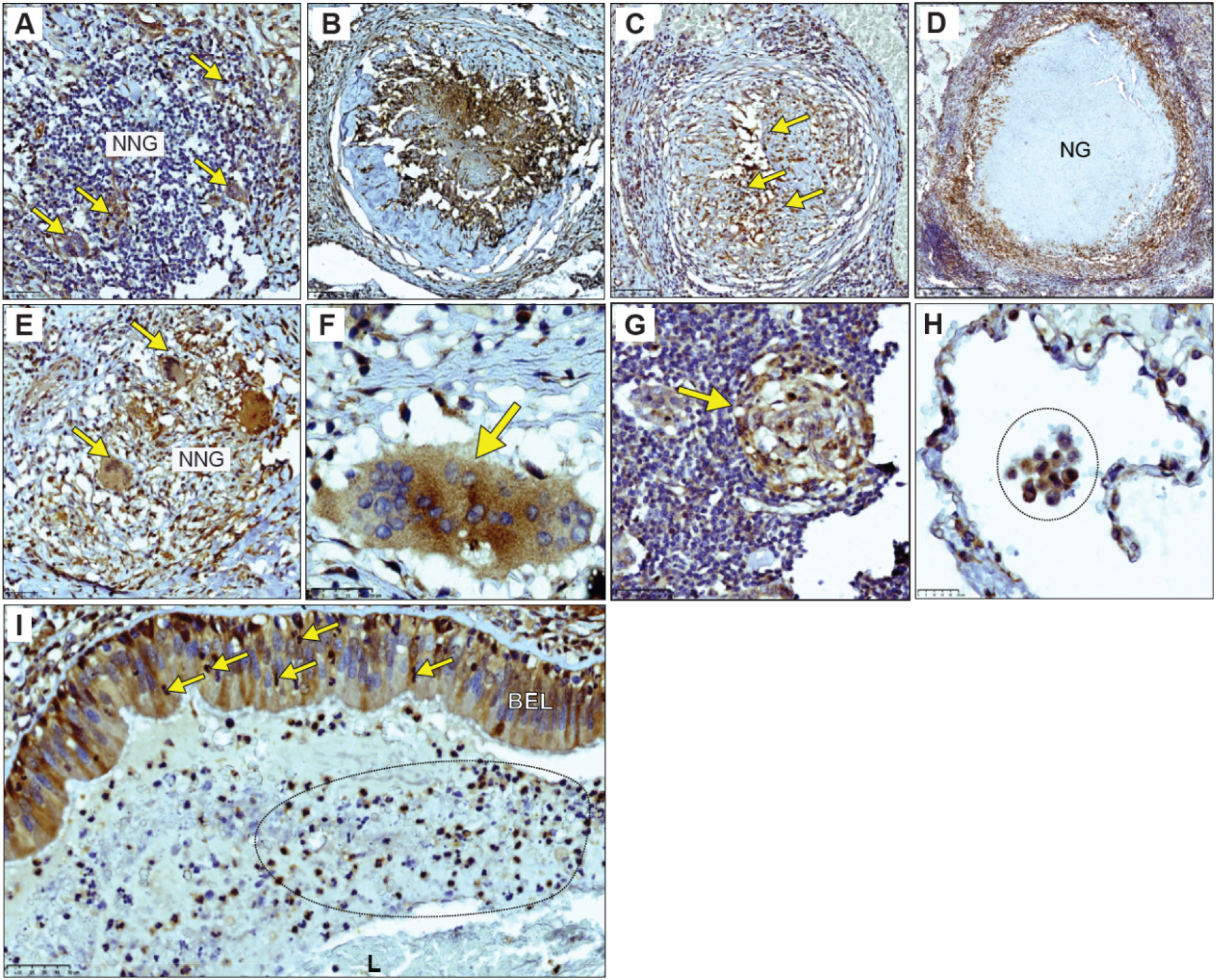
LDHA immunostaining across the histopathological spectrum of human TB. **(A)** Focal staining of LDHA (yellow arrows) within a non-necrotic granuloma (NNG). **(B)** Homogenous staining of LDHA within the necrotic center of an early necrotic granuloma. **(C)** Heterogeneous staining of LDHA within the necrotic center of a more developed necrotic lesion. Yellow arrows depict pyknotic nuclei within the necrotic center. **(D)** Staining of LDHA in the granulation layer and surrounding granulomatous inflammation region of a necrotic granuloma (NG) in the human TB lung. **(E)** Medium- and **(F)** high-magnification images of giant cells immunostained for LDHA (yellow arrows) in the context of **(E)** a non-necrotizing granuloma (NNG) and **(F)** necrotizing granuloma (NG). **(G)** LDHA staining of a lymphoid aggregate (yellow arrow) in a region proximal to a NG. **(H)** High-magnification image of LDHA staining in leukocytes (circled region) in an alveolus of a human TB patient. **(I)** LDHA staining in neutrophils (yellow arrows) crossing the positively-stained bronchial epithelial layer (BEL) in the context of inflammatory and necrotic debris (marked oval region) within the airway lumen (L).

Further association of LDHA with immune cell function in the human TB lung can be made based on positive LDHA staining within giant cells (Figure 1E, F; Figure S1A), lymphoid aggregates (Figure 1G), and alveolitis (Figure 1H; Figure S1B, C). The lack of LDHA staining in the immediate vicinity of these cells suggests that LDHA is linked to immune cell function as opposed to strictly a response to the surrounding microenvironment. LDHA is not expressed exclusively by immune cells in the human TB lung. We also observed positive LDHA staining in bronchial epithelial cells (Figure 1I; Figure S1D) and in the pulmonary vasculature (Figure S1E, F). Of note, we observed the LDHA-positive neutrophils crossing the bronchial epithelium and LDHA-positive neutrophils and macrophages among the inflammatory debris within the airway, underscoring a potential role for LDHA in the function of these cells (Figure 1I).

Taken together, histopathological appraisal of TB lesions provides new insight into the spatial distribution of LDHA within the human tuberculous lung. The distinct patterned responses within the spectrum of lesions were illustrated by myeloid, bronchial epithelial cells, and lymphocytes that stain positive for LDHA while engaging in distinctive immunological phenomena like granuloma formation and alveolitis. These data implicate LDHA as an important metabolic protein in the immune response in human TB lesions.

### Glycolytic capacity in myeloid cells protects mice against *Mtb* infection

Since NAD(H) regulates glycolysis at defined steps and the role of LDHA in TB pathogenesis is unknown, we hypothesized that NAD(H)-mediated glycolytic flux in myeloid cells protects the host against *Mtb* infection (Figure 2A, B). We tested this hypothesis using *Ldha*^*LysM*−/−^ mice which lack LDHA in the myeloid compartment^30^ (Figure 2C, D). Myeloid cells in these mice exhibit reduced LDH function as evidenced by a decreased ability to regenerate NAD^+^ from NADH in the presence of pyruvate (Figure 2E), which consequently reduces their glycolytic capacity (Figure 2F, G). We infected *Ldha*^*LysM*−/−^ and *Ldha*^fl/fl^ (WT) mice with a low-dose (∼30 colony forming units [CFU]) of *Mtb* H37Rv to assess their survival in a chronic model of TB. *Ldha*^*LysM*−/−^ mice were more susceptible to *Mtb* infection with significantly reduced survival (Figure 2H). We assessed the burden of *Mtb* and the pathology in the lungs and spleen of similarly low-dose infected mice at 4 weeks post infection (wpi), 10 wpi, and 30 wpi, which correspond to the early, middle, and late stages of the chronic TB model, respectively. At 4 wpi and 10 wpi, the *Mtb* burden in the lungs of *Ldha*^*LysM*−/−^ mice was significantly increased compared to WT mice. However, there was no difference in lung burden at 30 wpi (Figure 2I). The *Mtb* burden in the spleens of *Ldha*^*LysM*−/−^ mice was marginally increased at 4 wpi compared to WT, but no significant differences in *Mtb* burden were found at any time point (Figure S2A).

**Figure 2.**
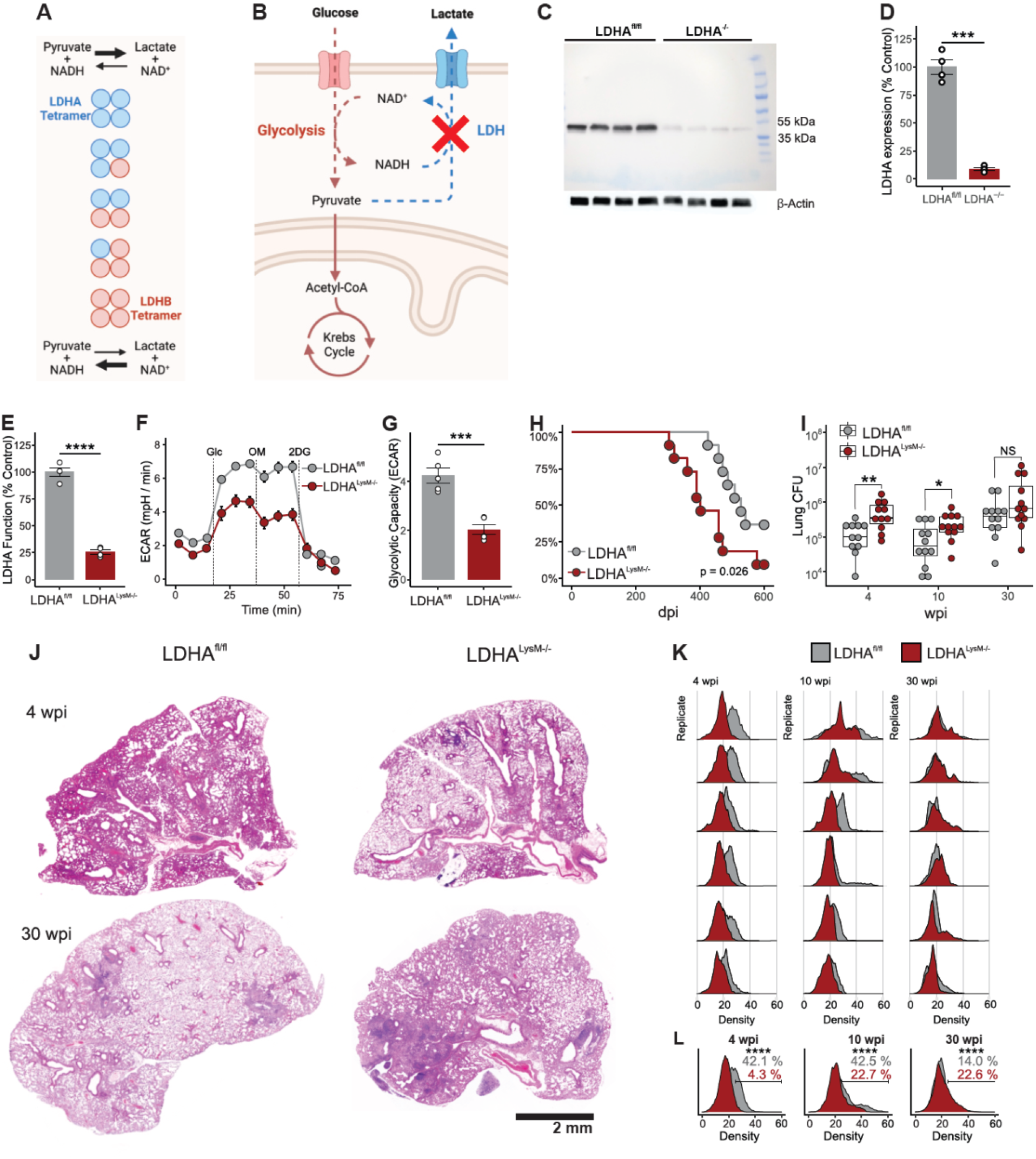
*Ldha*^*LysM−/−*^ mice are more susceptible to TB. **(A)** Summary figure depicting the preferential conversion of pyruvate to lactate by LDH composed predominantly of LDHA subunits compared to the preferential catalysis of the reverse reaction by LDH composed predominantly of LDHB subunits. **(B)** Summary figure depicting the glycolytic defect imposed by the deletion of LDHA within cells. **(C)** Immunoblot showing relative LDHA protein levels in LDHA^−/−^ and LDHA^fl/fl^ BMDMs. **(D**) LDHA protein levels and **(E)** LDH function measured in protein lysates and normalized to the mean values for LDHA^fl/fl^ BMDMs. **(F)** Line graph of the ECAR of LDHA^fl/fl^ and LDHA^−/−^ BMDMs. Dashed lines in represent the injection of glucose, oligomycin, and 2-DG, respectively. Symbols and error bars represent mean ± SEM of 5 technical replicates. **(G)** Columns, error bars, and symbols representing the mean, SEM, and individual values for glycolytic capacity determined from **(F)**. (**H)** Kaplan-Meier plot of mice infected with ∼25 colony forming units of *Mtb* (n = 11/group). **(I)** Box plot representing the burden of *Mtb* in the lungs of mice. Symbols represent biological replicates pooled from two independent experiments (n = 10 - 12/group). **(J)** Hematoxylin and eosin staining of representative lung sections collected from *Mtb* infected mice within the indicated conditions at the indicated time points. Scale bar represents 2 mm. **(K)** Ridgeline plot representing the normalized histograms for the cellular density around each nucleus in a tissue section. Shift of histograms to the right represents increased density of cells within a tissue section. Within each timepoint, biological replicates were ranked by mean density and plotted with the replicate of the corresponding rank from the other genotype. **(L)** Cumulative histogram of all the cells within a given genotype at a given time point compared to the other genotype within the same time point. Data are presented in **(D)** and **(E)** as individual values with the group mean ± SEM. Statistical significance was determined by two-sample t-test not assuming equal variance **(D, E, G)**, log-rank test **(H)**, two-sample Wilcoxon rank-sum test **(I)**, and two-sample z-test **(L)**. * p < 0.05, ** p < 0.01, *** p < 0.001, **** p < 0.0001.

Consistent with overall survival, qualitative assessment of the gross pathology (Figure S2B) and histopathology (Figure 2J) of the lungs revealed worse disease in *Ldha*^*LysM*−/−^ mice at 30 wpi. Interestingly, while lesions were well organized with a very high cell density at 30 wpi, inflammation was more diffuse at 4 wpi (Figure 2J). To accurately quantify this diffuse pathology, we took an unsupervised approach using the open-source application, QuPath^33^, to segment nuclei across each tissue section and determined the local density of cells for each nucleus. We found that WT mice initially mounted a robust inflammatory response that resolved by 30 wpi (Figure S2C), consistent with a more protective immune response to *Mtb* infection. We applied this approach to lungs from *Mtb*-infected *Ldha*^*LysM*−/−^ mice and found a striking absence of early inflammation in *Ldha*^*LysM−/−*^ mice compared to WT controls (Figure 2K, L; Figure S3). This deficit persisted through 10 wpi, and, while WT mice appeared to undergo significant resolution of inflammation at 30 wpi, inflammation in the lungs of *Ldha*^*LysM*−/−^ mice progressed through this time point (Figure 2K, L; Figure S3).

Altogether, we found that *Ldha*^*LysM*−/−^ mice are more susceptible to TB, maintain a higher *Mtb* burden in the lungs, and exhibit a dysfunctional inflammatory response to *Mtb* infection. This suggests that LDHA is necessary for protection against TB and that glycolytic flux in myeloid cells is essential for the control of *Mtb* infection and disease.

### Glycolytic myeloid cells are essential for protective immunity in TB

We observed reduced inflammation in the lungs of *Ldha*^*LysM−/−*^ mice at 4 wpi and increased inflammation at 30 wpi. To determine whether the dysfunctional immune response to *Mtb* infection was generalized across all leukocytes or unique to a particular subset or lineage, we used multiparameter flow cytometry to phenotype and quantify the immune cells trafficking to the lungs of *Ldha*^*LysM−/−*^ mice in our chronic model of TB (Figure S4A)^34^. The relative abundance of live, CD45+ cells (leukocytes) showed a trend consistent with our histology data, with *Ldha*^*LysM−/−*^ mice exhibiting decreased abundance of leukocytes at 4 wpi infection and increased abundance at 30 wpi, relative to WT controls (Figure 3A). The differences in leukocyte abundance at 0 wpi and 4 wpi arose from differences in abundance of several immune cell types (Figure 3B, S4B), but was limited to neutrophils and T cells at 10 wpi (Figure S4C) and T cells and monocytes at 30 wpi (Figure 3C). Of note, cell types belonging to both lymphoid and myeloid lineages were differentially abundant at all timepoints, indicating the origin of these differences lies in the recruitment of immune cells as opposed to differences in the trafficking or the viability of myeloid cells due to their glycolytic deficiency.

**Figure 3.**
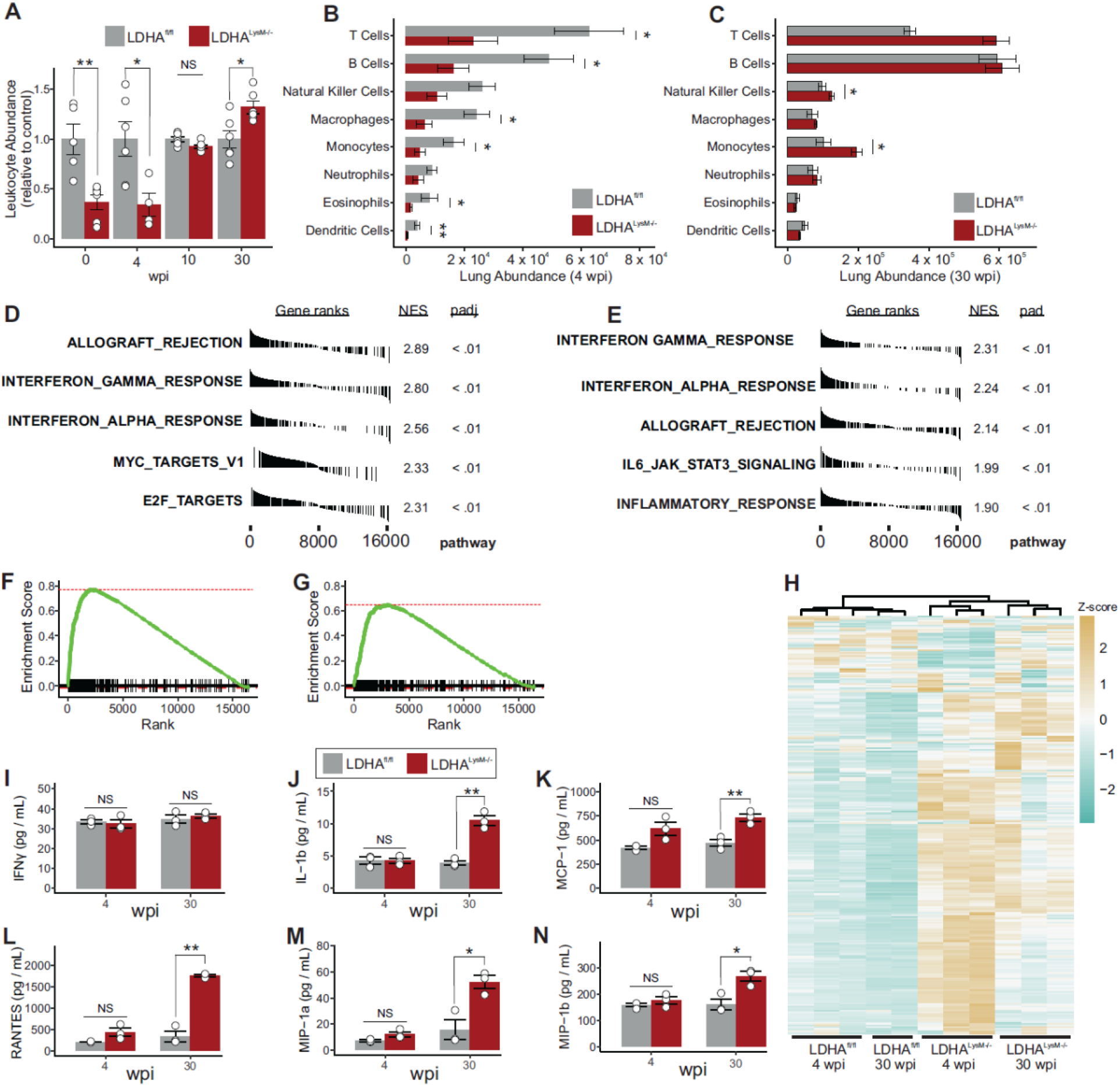
*Ldha*^*LysM−/−*^ mice exhibit a dysregulated immune response to *Mtb* infection. **(A)** Column graph representing the mean abundance of leukocytes within the lungs of mice relative to the mean abundance of leukocytes in WT mice at each time point (n = 4-6/group). **(B, C)** Bar graphs depicting the absolute count of the indicated immune cell population in the lungs of *Mtb*-infected mice at **(B)** 4 wpi and **(C)** 30 wpi (n=6/group). **(D-G)** Tables of the most enriched hallmark gene sets **(D, E)** and enrichment plots **(F, G)** for the transcriptional response to IFNγ stimulation in the lungs of mice at 4 wpi **(D, F)** and 30 wpi **(E, G)** with *Mtb*. **(H)** Heatmap representing the expression of the genes comprising the hallmark response to IFNγ gene set. Color-scale values represent the z-score for the expression of a given row and the dendrogram above the columns represents the results of hierarchical clustering based on the expression of these genes. **(I-N)** Column graphs representing the protein expression of the indicated cytokines in the lungs of *Mtb*-infected mice (n=3/group). Data are presented as individual values with the group mean ± SEM for panels **(A-C)** and **(I-N)**. Statistical significance was determined by the two-sample Wilcoxon rank-sum test **(A-C)** and two-sample t-test not assuming equal variance. * p < 0.05, ** p < 0.01.

To better understand functional aspects of the immune response in the lungs of *Ldha*^*LysM−/−*^ mice, we performed RNA sequencing (RNA-Seq) in the lungs of these mice at 4 wpi and 30 wpi. To assess global differences in lung transcriptomes, we performed principal component analysis, which separates replicates by group across the first two components (Figure S4D). To identify the biological processes that best distinguished these groups, we analyzed differentially expressed genes using gene set enrichment analysis (GSEA) to identify crucial biological processes for which transcripts are enriched or deficient in *Ldha*^*LysM−/−*^ mice^35^. Seemingly contrary to our evidence of a reduced immune response at 4 wpi, transcripts associated with inflammatory processes were among the most enriched transcripts in *Ldha*^*LysM−/−*^ mice at 4 wpi and 30 wpi among the 50 hallmark gene sets curated for GSEA (Figure 3D, E). In particular, the response to IFNγ was the most enriched gene set at both time points (Figure 3D-G). This relationship was further underscored by the ability of hierarchical clustering to separate the groups of samples based on their expression of the genes in this gene set (Figure 3H). The robust IFNγ gene signature in more susceptible mice with a blunted immune response is particularly intriguing since IFNγ is an indispensable antimycobacterial cytokine widely considered to be protective in TB^36,37^.

To determine whether the enrichment of mRNA corresponded to increased levels of cytokines and chemokines, we performed a multiplexed, fluorescence-based assay to quantify the expression of cytokines and chemokines in lung tissue from the same mice for which sequencing data was available. Despite an increase in transcription, we observed similar levels of nearly all cytokines and chemokines between WT and KO mice at 4 wpi, including IFNγ (Figure 3I-N; Figure S4E-G). However, at 30 wpi levels of several key cytokines and chemokines were increased in *Ldha*^*LysM*−/−^ animals compared to WT controls, consistent with other signs of chronic inflammation (Figure 3J-N; Figure S4E-G). Consistent with our flow cytometry data, we observed increased concentrations of chemokines associated with the recruitment of monocytes (MCP-1, KC), T cells (RANTES), or both (MIP-1a, MIP-1b).

Taken together, these results indicate glycolytic myeloid cells are important for the early recruitment of both lymphocytes and other myeloid cells in response to *Mtb* infection. Furthermore, given the discrepancy between having a robust IFNγ transcriptional signature and increased bacterial burden in the lungs of *Ldha*^*LysM−/−*^ mice, our data complement previous studies^15,18,22^ by suggesting that glycolytic myeloid cells are essential mediators of the protective effects of IFNγ *in vivo*.

### Macrophages require LDHA for their metabolic response to IFNγ

Based on our RNA-Seq analysis showing increased IFNγ signaling in the context of poorly controlled *Mtb* infection in *Ldha*^*LysM−/−*^ mice, we next sought to understand the role of lactate metabolism in the myeloid response to IFNγ. We hypothesized that LDHA-deficient macrophages are unable to respond metabolically to IFNγ. Therefore, we performed combined glycolysis/mitochondrial stress tests on bone marrow-derived macrophages (BMDMs) as reported elsewhere^38^. Because this technique determines the non-glycolytic acidification caused by cells based on the extracellular acidification rate (ECAR) prior to the injection of glucose rather than after the injection of 2DG, it is important to validate that these values are the same in a standard glycolysis stress test. Indeed, we found no difference in these two approaches in a standard glycolysis stress test (Figure S5A, B). Consistent with our hypothesis, we observed only a modest difference in glycolytic capacity between LDHA^−/−^ and WT BMDMs without IFNγ (Figure 4A). However, upon stimulation with IFNγ, the increase in glycolytic capacity for LDHA^−/−^ BMDMs was threefold lower than in WT BMDMs (Figure 4A, B; Figure S5C-E). Furthermore, while mitochondrial respiration in LDHA^−/−^ BMDMs was unchanged by stimulation with IFNγ (Figure 4C, D), these cells were more reliant on OXPHOS at baseline (Figure S5G-J) and had reduced spare respiratory capacity (SRC) (Figure S5K). Taken together, these data demonstrate that LDHA^−/−^ macrophages exhibit no bioenergetic reserve (Figure 4E, F), hampering their metabolic response to IFNγ.

**Figure 4.**
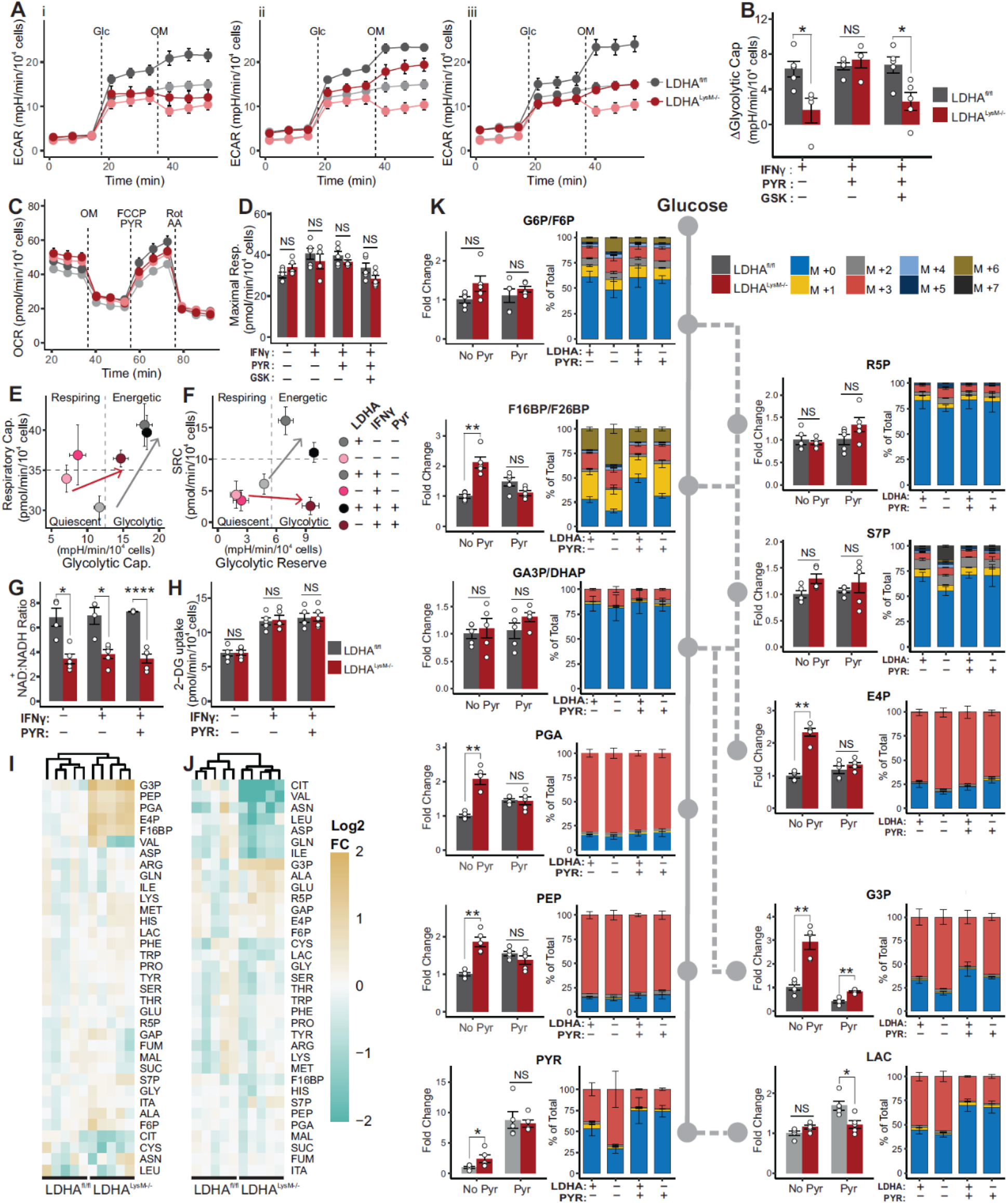
LDHA^−/−^ BMDMs exhibit pyruvate-reversible deficiency in their metabolic response to IFNγ. **(A)** Line graphs of the ECAR of LDHA^−/−^ (red) and WT (gray) BMDMs treated with (i) IFNγ (10 ng/mL); (ii) IFNγ and 1 mM pyruvate; and (iii) IFNγ, pyruvate, and 5 μM GSK 2837808 (darker lines and symbols) relative to untreated BMDMs (lighter lines and symbols). Dashed lines in each panel represent an injection of glucose (Glc) followed by an injection of oligomycin (OM). Symbols and error bars represent mean ± SEM of 5 biological replicates. **(B)** Column graph representing the change in glycolytic capacity of BMDMs treated as indicated in (**A**) compared to unstimulated BMDMs (n = 5/group). **(C)** Line graph of the OCR of BMDMs treated with 10 ng/mL IFNγ (darker lines and symbols) or no stimulus (lighter lines and symbols). Dashed lines in each panel represent the following injections: OM, FCCP/Pyruvate, and Rotenone/Antimycin A. Symbols and error bars represent mean ± SEM of 5 biological replicates. **(D)** Column graph representing the maximal respiration of BMDMs treated as indicated (n = 5/group). **(E, F)** Phenograms for the indicated parameters extracted from the XF profiles of BMDMs treated with the indicated conditions. Symbols and error bars represent the mean ± SEM for 5 biological replicates. **(G, H)** Column graphs representing G) the NAD^+^:NADH ratio or H) 2DG uptake of BMDMs treated as indicated (n = 4/group). **(I, J)** Heatmaps representing the abundance of the metabolites comprising key pathways of central carbon metabolism in BMDMs treated with I) 10 ng / mL IFNγ alone or J) 10 ng/mL IFNγ and 1 mM pyruvate. Color-scale values represent the Log_2_ fold-change relative to the mean value for WT BMDMs. The dendrogram above the columns represents hierarchical clustering of replicates based on the abundance of these metabolites. **(K)** Column graphs of the abundance (left) and isotopologue distribution (right) of the indicated metabolite and its position within glycolysis (solid line), the PPP (dashed line), and related pathways (dotted lines). Column graphs (left) represent the fold change (FC) in abundance of the indicated metabolite relative to the mean of WT BMDMs stimulated with IFNγ. Stacked column graphs (right) corresponding to 100% of the total abundance for each metabolite, where the proportion of each mass isotopologue for the indicated metabolite is indicated by color. Data are presented as individual values for biological replicates with the group mean ± SEM for panels **(B, D, G, H, and K)**. Statistical significance was determined by two-sample t-test without assuming equal variance **(B, D, G, H, K)**. * *p* < 0.05, ** *p* < 0.01, *** *p* < 0.001, **** *p* < 0.0001.

In our initial combined stress tests, injection of FCCP and pyruvate elicited a marked increase in ECAR in LDHA^−/−^ BMDMs (Figure S5F). We posited that supplementing LDHA^−/−^ BMDMs with pyruvate would increase lactate fermentation catalyzed by LDH composed of LDHB subunits, with a concomitant increase in limiting NAD^+^. We found that while pyruvate supplementation did little to impact basal glycolytic flux (Figure S5D), it rescued the deficit in the glycolytic response to IFNγ (Figure 4A, B) and the deficit in glycolytic reserve in LDHA^−/−^ BMDMs (Figure S5E), thereby reducing their deficit in glycolytic capacity (Figure S5C). Importantly, this rescue was attenuated by the selective LDH inhibitor GSK 2837808A, indicating the effect is mediated by residual LDH function (*i*.*e*., LDHB) in LDHA^−/−^ BMDMs (Figure 4A, B; Figure S5C-E).

To further characterize the glycolytic metabolism of LDHA^−/−^ BMDMs, we assessed the NAD^+^:NADH ratio and glucose uptake in these cells following stimulation with IFNγ and pyruvate supplementation. As expected, LDHA^−/−^ BMDMs exhibited a decreased NAD^+^:NADH ratio regardless of treatment condition (Figure 4G). Interestingly, however, glucose uptake was no different between LDHA^−/−^ and WT BMDMs (Figure 4H). Considering the comparable glucose uptake but diminished metabolic flux through glycolysis, we hypothesize that IFNγ-stimulated LDHA^−/−^ BMDMs would accumulate metabolic intermediates within glycolysis and the reversible reactions of the non-oxidative PPP. Therefore, we performed targeted metabolomics with uniformly labeled ^13^C_6_ glucose, which revealed a cluster of metabolites (F16BP, PGA, PEP, E4P, and G3P) with significantly increased abundance in LDHA^−/−^ BMDMs which falls between the rate-limiting steps in upper and lower glycolysis and the irreversible oxidative PPP (Figure 4I). Consistent with our bioenergetics analysis, pyruvate supplementation abrogated this accumulation (Figure 4J, K; Figure S5O). Interestingly, while the observed cluster of glycolytic and PPP metabolites was no longer present in pyruvate-supplemented BMDMs, metabolites in a new cluster were maintained at reduced abundance in LDHA^−/−^ BMDMs (Figure 4J). This new cluster is comprised of important TCA cycle intermediates, such as citrate, as well as amino acids *(e*.*g*., Val, Asn, Leu, Asp, Gln, Ile) that are carbon sinks for TCA cycle intermediates. These findings are consistent with excess pyruvate being utilized by WT BMDMs as a carbon source for the TCA cycle, whereas LDHA^−/−^ BMDMs primarily rely on pyruvate as an electron sink to regenerate NAD^+^.

In summary, these data suggest that LDH-mediated NAD^+^ regeneration is essential for the metabolic response to IFNγ. Of note, we also found that pyruvate supplementation can have the counterintuitive effect of facilitating glycolytic flux in the setting of reduced LDH enzymatic capacity, opening the door for a new approach to the enhancement of glycolysis under appropriate conditions.

### *Mtb* blunts the metabolic response to IFNγ in macrophages

Our data show that glycolysis in myeloid cells, which depends strictly on NAD^+^ availability, is essential for protective immunity mediated by IFNγ. Given that virulent *Mtb* also reduces glycolytic flux in macrophages relative to killed or attenuated *Mtb*^16,19,20^, we next hypothesized that, like *LDHA* deletion, virulent *Mtb* blunts the metabolic response to IFNγ in infected macrophages. Consistent with our hypothesis, *Mtb* infection increased the glycolytic capacity of WT BMDMs relative to no stimulus. However, these cells were unable to increase their glycolytic capacity in response to IFNγ stimulation compared to uninfected BMDMs (Figure 5A, B). Unlike LDHA^−/−^ BMDMs, however, the blunted response to IFNγ induced by *Mtb* infection could not be rescued by pyruvate supplementation (Figure S6A, B), suggesting a mechanism distinct from reduced LDH expression. We therefore performed targeted metabolomics to characterize the central carbon metabolism in *Mtb*-infected BMDMs with or without IFNγ. We found *Mtb* infection decreased the abundance of glycolytic and PPP intermediates in BMDMs, except for pyruvate, which was significantly increased compared to uninfected macrophages with or without IFNγ-stimulation (Figure 5C). In addition, analysis of TCA cycle intermediates revealed a comparable increase in the pool of (iso)citrate in *Mtb*-infected BMDMs. These data are consistent with a mechanism whereby the capacity of macrophages to funnel pyruvate into lactate metabolism is limited, leading to an accumulation of pyruvate, and subsequently, (iso)citrate, which then allosterically inhibits the rate-limiting enzyme of glycolysis, phosphofructokinase, further reducing glycolytic flux^39^.

**Figure 5.**
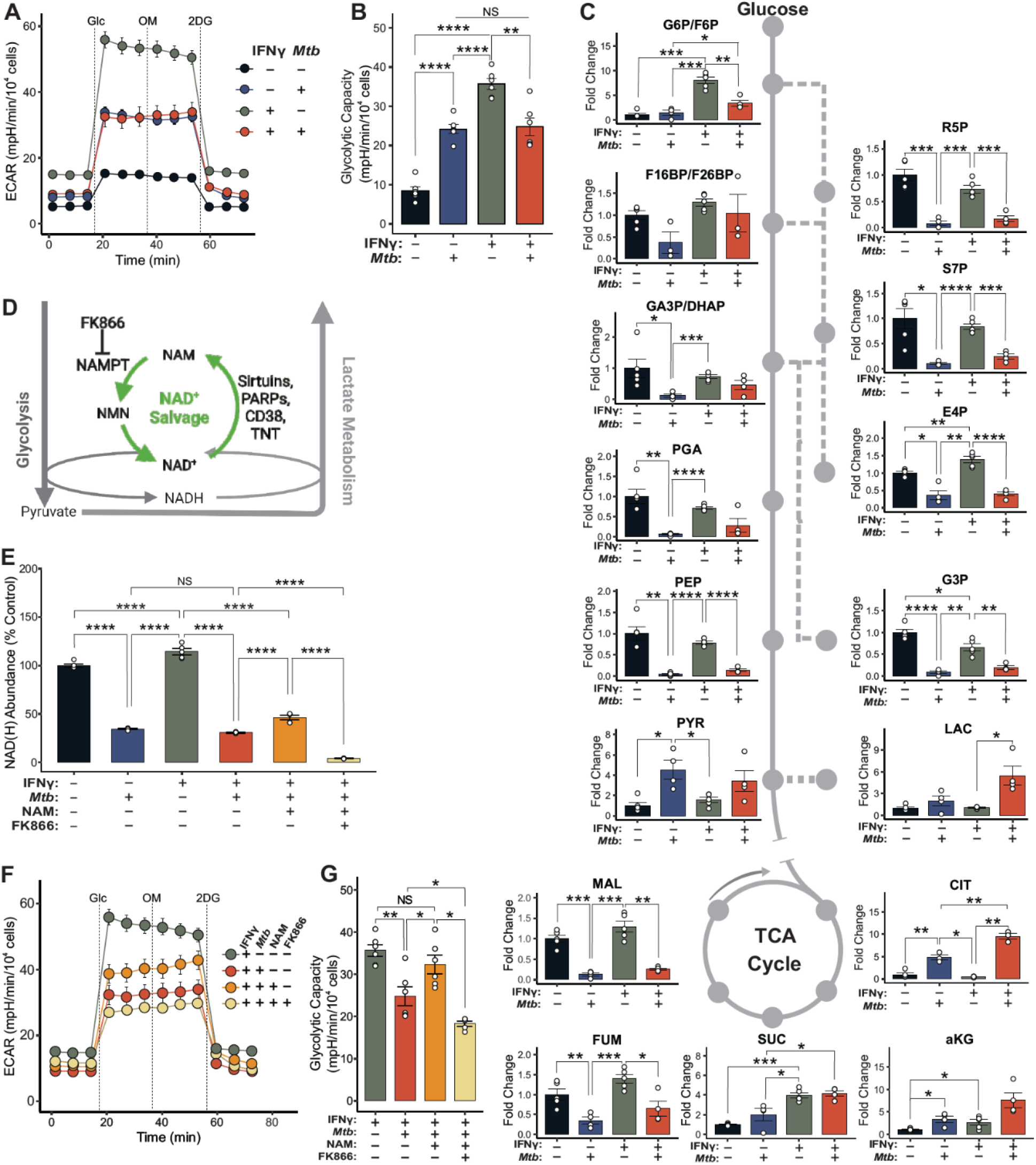
*Mtb* disrupts glycolysis by reducing NAD(H) abundance in infected macrophages. **(A)** ECAR profiles for WT BMDMs exposed to combinations of IFNγ (10 ng/mL) and *Mtb* (MOI 5:1) for 18 hours. Dashed lines represent injections of glucose (Glc), oligomycin (OM), and 2DG, respectively. Symbols and error bars represent mean ± SEM of 6 biological replicates. **(B)** Column graph representing the glycolytic capacity determined from the profiles in panel A (n = 6/group). **(C)** Column graphs of the abundance of the indicated metabolite and its position within glycolysis (solid line), the PPP (dashed line), the TCA cycle (solid circle), and related pathways (dotted lines) relative to the mean of uninfected, unstimulated LDHA^fl/fl^ BMDMs. Values for IFNγ-treated BMDMs are repeated from Figure 4K. **(D)** The NAD^+^ salvage pathway (green), key enzymes, and the inhibitor, FK866, and its relation to glycolysis and lactate metabolism. **(E)** Column graph representing the total abundance of NAD(H) within BMDMs under the indicated conditions (n = 5/group). **(F)** ECAR profiles for BMDMs exposed to combinations of IFNγ (10 ng/mL), *Mtb* (MOI 5:1), NAM (1 mM) and FK866 (200 nM) for 18 hours. Dashed lines represent injections of Glc, OM, and 2DG, respectively. Symbols and error bars represent mean ± SEM of 6 biological replicates. Values for IFNγ and IFNγ + *Mtb* from **A. (G)** Column graph representing the glycolytic capacity determined from the profiles in panel A (n = 6/group). Values for IFNγ and IFNγ + *Mtb* are repeated from **B**. Data are presented as individual values for biological replicates with the group mean ± SEM for panels **(B, C, and G)** and as individual values for technical replicates with the group mean ± SEM for panel **(E)**. Statistical significance was determined by two-sample t-test without assuming equal variance (B, C, E, G). * *p* < 0.05, ** *p* < 0.01, *** *p* < 0.001, **** *p* < 0.0001.

Together, these data suggest that while *Mtb*-infected macrophages likely have the enzymatic capacity to metabolize excess pyruvate to lactate, they are unable to do so. We then considered the contribution of NADH, given that it is the only other factor required for this reaction. *Mtb* was shown to reduce NAD(H) levels in infected macrophages^40,41^, and our own bioenergetic and metabolomic analyses are consistent with the metabolic profile of inflammatory macrophages following inhibition of the NAD^+^ salvage pathway (Figure 5D)^25^. Based on these considerations, we hypothesize that glycolysis is inhibited in *Mtb*-infected BMDMs due primarily to a reduction in NAD(H) availability. Consistent with our hypothesis, while IFNγ stimulation increases the NAD(H) pool in macrophages, *Mtb* infection reduces the abundance of NAD(H) whether or not IFNγ is present (Figure 5E). This decrease could be partially rescued by exposing macrophages to nicotinamide (NAM), which can be converted to NAD(H) via the NAD^+^ salvage pathway. This rescue, as well as the residual NAD(H) pool in these cells, were eliminated by FK866, an inhibitor of nicotinamide phosphoribosyltransferase (NAMPT) the rate-limiting enzyme in NAD^+^ salvage (Figure 5E). To determine if the decreased glycolytic capacity in these cells resulted from reduced NAD(H) levels, we performed bioenergetic analysis on *Mtb*-infected BMDMs supplemented with NAM. Addition of NAM restored the glycolytic capacity of *Mtb*-infected BMDMs and FK866 blocked this effect (Figure 5F, G). Consistent with FK866-mediated reduction in NAD(H) levels, treatment with FK866 further reduced glycolytic capacity below that of *Mtb*-infected, IFNγ-stimulated BMDMs. This suggests that NAD^+^ salvage is partially responsible for the maintenance of glycolytic capacity following infection with *Mtb*, even in the absence of NAM.

In summary, these data demonstrate that *Mtb* reduces NAD(H) availability, and that NAM restores the glycolytic capacity of *Mtb*-infected BMDMs by restoring NAD(H) levels through its conversion to NAD(H) via the NAD^+^ salvage pathway. Furthermore, these findings underscore the importance of mechanistic insight into the pathogenesis of *Mtb*, as pyruvate supplementation, which restored the glycolytic capacity of *LDHA*^−/−^ macrophages (Figure 4B), was ineffective at restoring the glycolytic capacity of WT macrophages infected with *Mtb* (Figure S6A, B).

### Nicotinamide is an effective treatment for TB

NAM is known to be an effective treatment for TB^42–44^, and was recently shown to exert an uncharacterized, primarily host-directed effect^45^. Based on our observations that NAM increases the glycolytic capacity of *Mtb*-infected BMDMs and that glycolysis in myeloid cells promotes bacterial control *in vivo*, we hypothesize that NAM acts as a host-directed therapy (HDT) by enhancing glycolysis in *Mtb*-infected macrophages through its conversion to NAD(H) via the NAD^+^ salvage pathway. Therefore, we infected BMDMs with luciferase-expressing *Mtb* and measured luminescence at 48 hpi across a range of NAM conditions. We observed that NAM exerts a dose-dependent reduction in bacterial luminescence signal in BMDMs (Figure 6A). Consistent with previous work^45^, exposure to NAM modestly reduced *Mtb* growth in 7H9 broth, indicating that the therapeutic effect of NAM is host-directed. We next performed this experiment in the presence of NAM with addition of FK866 or 2DG. In both cases, we observed that the antimycobacterial effect of NAM, but not the pathogen-directed antimycobacterial drug, isoniazid, was reversed by the respective inhibitor (Figure 6B, C). This indicates that NAM exerts a host-directed effect in macrophages by enhancing glycolytic flux through its conversion to NAD^+^. Additionally, FK866 and 2DG significantly increased the bacterial burden in the absence of any treatment, further indicating that BMDMs rely on the NAD^+^ salvage pathway at baseline to maintain antimycobacterial function and that glycolysis is required for host protection.

**Figure 6.**
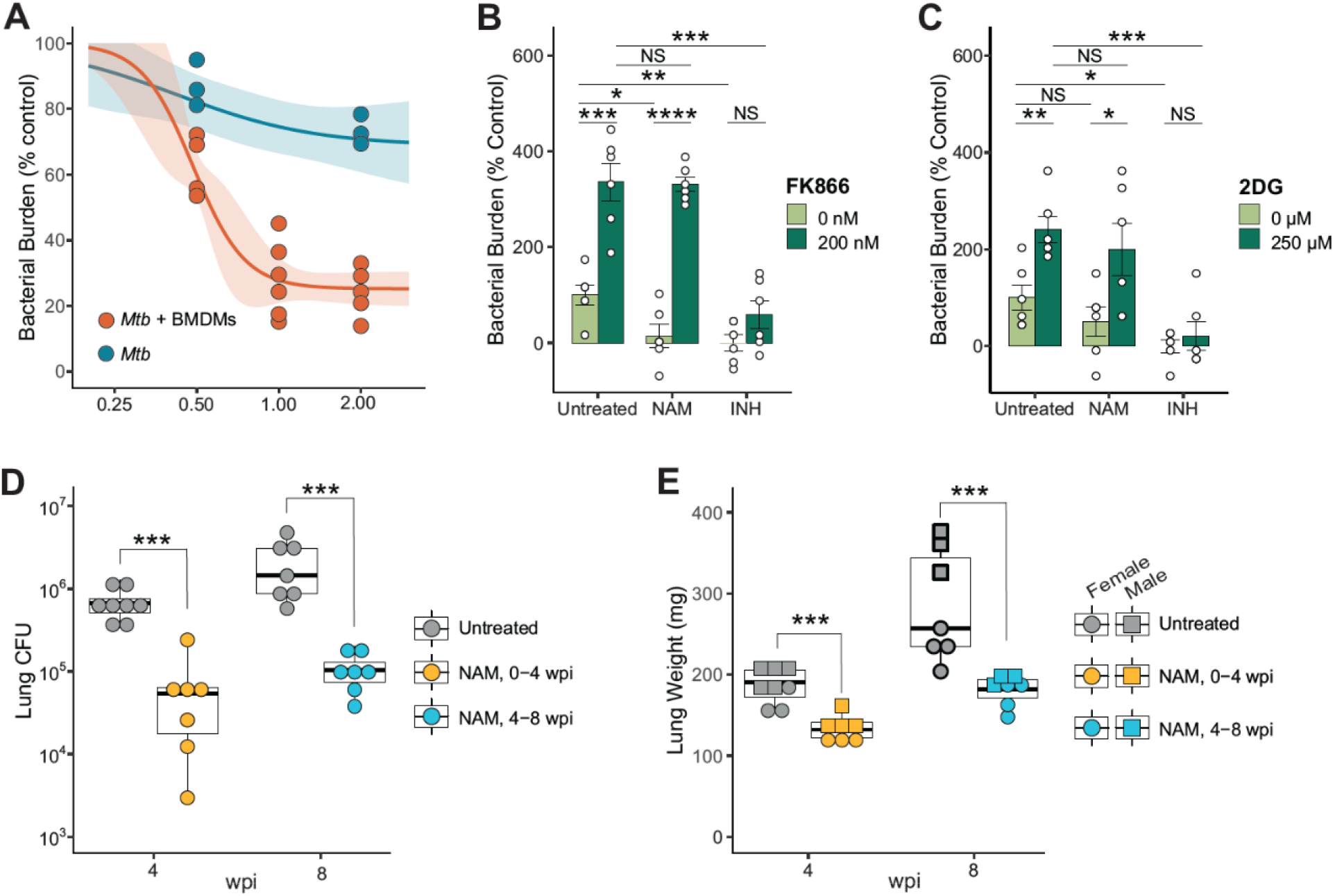
Nicotinamide is an effective host-directed therapy for TB. **(A)** Dose response curves for the antimycobacterial effect of NAM in 7H9 broth (blue) or infected BMDMs (orange). The shaded region around each curve represents the 95 % confidence interval, while dots represent biological replicates tested across the indicated concentrations. **(B, C)** Column graphs representing the bacterial burden in BMDMs infected with *Mtb* expressing a luciferase reporter treated with a combination of NAM and FK866 **(B)** or NAM and 2-DG **(C)** (n = 6/group). Isoniazid (INH) represents a direct-acting antimycobacterial compound, included as a control. Data are shown as a percentage of the mean value of the untreated, uninfected group. **(D, E)** Box plot representing the lung burden of *Mtb* **(D)** and lung weight **(E)** in mice treated with NAM for four weeks, beginning either at 3 days post infection of 4 weeks post infection. Symbols represent biological replicates (n = 7-8/group). Data are presented as individual values for biological replicates with the group mean ± SEM for panels **(B and C)**. Statistical significance was determined by two-sample t-test not assuming equal variance **(B, C, E)** or two-sample Wilcoxon rank-sum test **(D)**. * p < 0.05, ** p < 0.01, *** p < 0.001, **** p < 0.0001.

Finally, we evaluated the suitability of NAM as a treatment for TB in the mouse model of TB. Mice were infected (∼100 CFU) via the aerosol route and treated for four weeks starting at either 3 dpi (post-exposure prophylaxis) or 28 dpi (treatment). Both prophylactic and therapeutic approaches with NAM resulted in a statistically significant, ten-fold reduction in the bacterial burden in the lungs of mice (Figure 6D), as well as a significant decrease in lung weight reflecting reduced inflammation (Figure 6E).

In summary, our *in vitro* data show that NAM enhances macrophage control of *Mtb* infection specifically by its conversion to NAD^+^ through the NAD^+^ salvage pathway and its subsequent potentiation of glycolysis. Notably, our *in vivo* studies illustrate that orally administered NAM is effective as prophylaxis and treatment for *Mtb* infection. Overall, our findings point to NAM as a potential HDT for the treatment of TB.

## Discussion

The major conclusion of this study is that NAD(H) homeostasis in myeloid cells, maintained by LDHA activity and NAD^+^ salvage, supports glycolytic capacity and, subsequently, host protection in TB (Figure 7). Importantly, the clinical relevance of this work is rooted in the spatial and cellular distribution of LDHA in the spectrum of human TB lesions. Our *in vivo* experiments subsequently demonstrated that maximal glycolytic capacity in myeloid cells supports the induction of protective immunity, likely through the antimycobacterial effects of IFNγ. Our bioenergetic and pharmacological inhibition experiments show that *Mtb* disrupts glycolytic flux in infected macrophages by reducing the total abundance of the NAD(H) pool. We demonstrate that this virulence strategy can be countered through oral administration of the NAD^+^ precursor, NAM, constituting an effective HDT for TB. Our study represents a significant advance over previous work in the TB field that relies solely on 2DG or disruption of *Hif1A*, which not only suppress glycolysis, but also several other essential pathways. Overall, our findings provide fundamental new insight into how *Mtb* reprograms host glycolysis and highlight this pathway as a potential therapeutic target to control TB disease.

**Figure 7.**
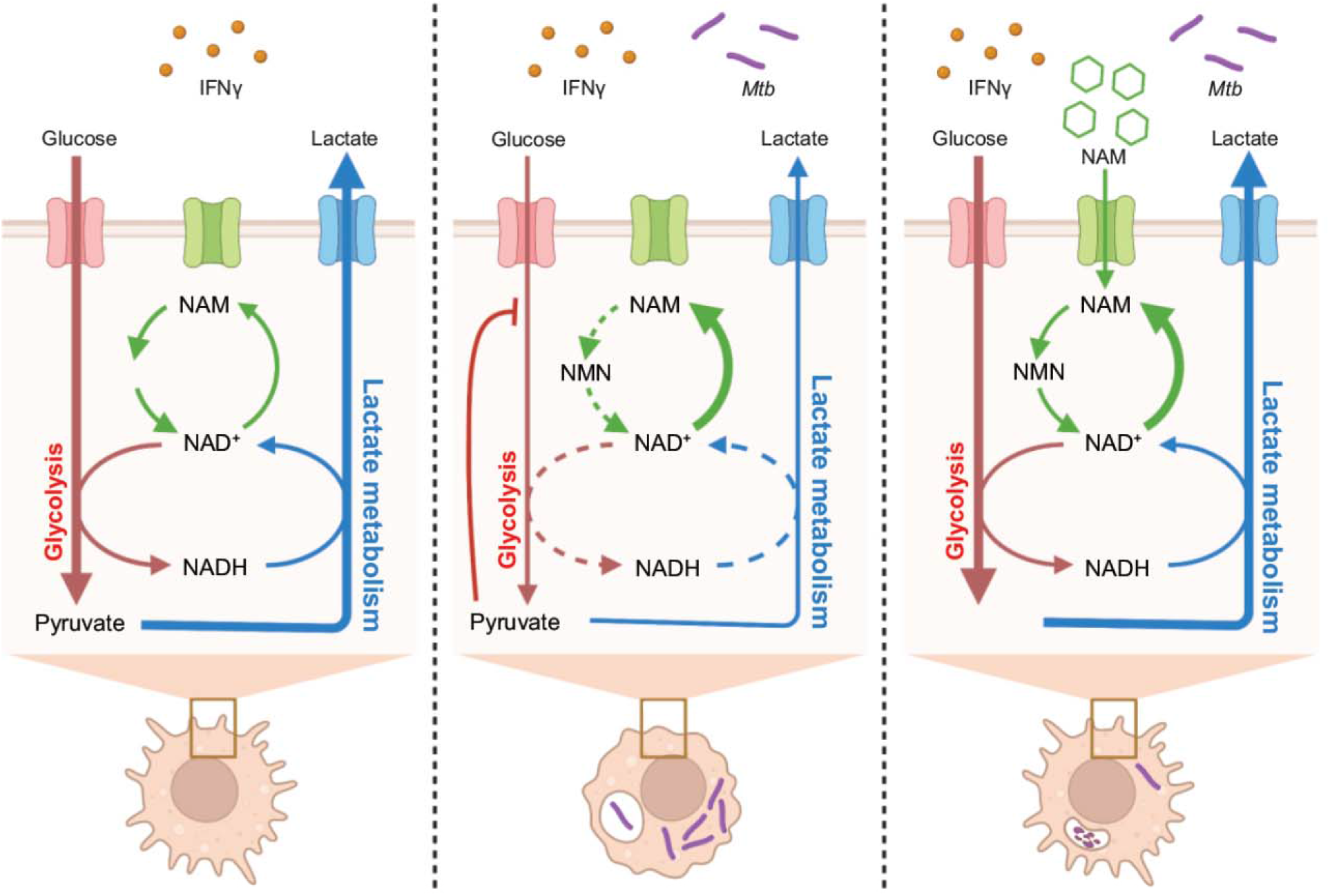
Nicotinamide protects the host from *Mtb* by maintaining NAD(H) homeostasis. Summary graphic depicting the proposed model for the metabolism of macrophages responding to stimulation with IFNγ, how *Mtb* disrupts this response, and how NAM restores it. **(A)** Following stimulation with IFNγ, macrophages exhibit high rates of glycolysis and lactate metabolism, necessitating the rapid cycling between NAD+ and NADH. Under these conditions, the NAD+ salvage pathway effectively maintains the pool of NAD(H). Together, this metabolic response enables the functional activation of the macrophage by IFNγ. **(B)** In *Mtb*-infected macrophages, the metabolic response to IFNγ is prevented by disrupting NAD(H) homeostasis. This may be due to decreased synthesis or increased NAD+ consumption. Regardless, reduced NAD(H) availability impairs the conversion of pyruvate to lactate, leading to an accumulation of pyruvate. Pyruvate is then converted to citrate which inhibits glycolytic flux. The functional consequence is an impairment in the host-protective effects of IFNγ. **(C)** Following treatment with NAM, *Mtb*-infected macrophages can respond effectively to IFNγ, as the NAM is metabolized to NAD+. While the insult to NAD(H) homeostasis is likely still present, NAM supplementation addresses the immediate downstream impact of this insult by supporting NAD+ synthesis. Consequently, the metabolic and functional responses to IFNγ and other environmental signals is restored, ultimately leading to host protection in TB.

This study affirms our central hypothesis that glycolytic flux in myeloid cells is essential for host protection in TB. Importantly, the use of *Ldha*^*LysM−/−*^ mice extends previous studies that associate glycolytic myeloid cells and host protection by applying a highly selective approach to the inhibition of glycolytic flux specifically in this subset of immune cells, rather than using an inhibitor of glucose uptake, 2DG^15,17,18^, which also suppresses the PPP and TCA cycle. Since glycolysis requires NAD^+^ for the oxidation of glyceraldehyde-3-phosphate to 1,3-bisphosphoglycerate, and NAD^+^ can be regenerated through the reduction of the glycolytic end product, pyruvate, by LDHA, the target of our enzymatic manipulation is external to process of glycolysis altogether. Hence, we effectively used this endogenous regulatory strategy for glycolysis, *i*.*e*., NAD^+^ availability, as a metabolic valve to precisely modulate glycolytic flux for the control of *Mtb* disease.

Conditional or whole-body knockouts of the transcription factor HIF1α have been used to examine the role of glycolysis during *Mtb* infection^15,18^; however, the resultant pleiotropic effects may confound interpretation of results. Thus, it is clear that inhibiting glycolysis, and more specifically inhibiting LDHA, is not a suitable HDT approach for TB. Interest in this approach has been based largely on the association between glycolysis, inflammation, and TB disease^3,14,46^. However, our *in vivo* findings suggest that while reduced glycolytic capacity in myeloid cells may reduce inflammation in the short term, it ultimately leads to chronic inflammation, worse pathology, and worse outcomes. Our data may seem to conflict with a report that *Mtb*-infected mice treated with FX11, a NADH-analog inhibitor of LDH, showed modest improvements in bacterial burden and pathology^47^. However, these results might be explained by a short-term reduction in inflammation, since the authors state that the effect of FX11 *in vivo* could not be confirmed as the on-target inhibition of LDH, leaving open the potential for other mechanisms for the protective effect.

Following our initial experiments, we limited our investigation in primary cells to the response to IFNγ, the most critical process as identified by our unsupervised analysis of 50 major cellular pathways and supported by the TB literature^15,18,19^. Similarly, we focused our studies on macrophages to best complement this literature, as well as studies of *Mtb*-induced glycolytic defects^16,19,20^, all of which center on macrophages. However, the reduction of glycolytic capacity due to NAD(H) depletion following *Mtb* infection is likely sufficiently generalizable to disrupt a wide array of glycolysis-dependent functions, including responses to pathogen-associated molecular patterns, damage-associated molecular patterns and other immunomodulatory signals^25^. This point is particularly well illustrated by the observation that heat-killed or irradiated *Mtb* induces a robust glycolytic response while virulent *Mtb* does not, *i*.*e*., while the physical components of the bacillus are sufficient to induce a robust glycolytic response, viable virulent *Mtb* suppresses that very response^16,19,48^. Our data in *Ldha*^*LysM−/−*^ mice suggest that this disruption of glycolysis may underlie the suppression of adaptive immunity in TB, as evidenced by the dramatically delayed immune response in these mice. Thus, per these observations and the extensive literature on immunometabolism, disruption of this central pathway likely has many consequences for the host.

Our findings raise the following question: how does *Mtb* deplete NAD(H) levels? Although outside the scope of the study, this effect, and why it is limited to virulent *Mtb*, can be partly explained by the secretion of tuberculosis necrotizing toxin (TNT), a NAD^+^ glycohydrolase^49^. WT *Mtb* was shown to significantly reduce NAD^+^ abundance in infected macrophages; however *Mtb* lacking functional TNT still reduced NAD^+^ to around 50% of WT *Mtb*^40,41^. This discrepancy appears to be explained by the observation that macrophages treated with lipopolysaccharide, a constituent of the outer membrane of gram-negative bacteria, exhibit a similar reduction in intracellular NAD(H) and increased reliance on the NAD^+^ salvage pathway^25^. In this context, glycolysis was not affected unless NAD^+^ salvage was also inhibited. Thus, NAD(H) depletion and inhibition of glycolysis may result from both an increased reliance on NAD^+^ salvage and increased strain on this pathway due to NAD^+^ degradation by TNT (Figure 7B).

Given the downstream consequences of NAD(H) depletion and inhibition of glycolysis, we attempted to address the upstream cause by applying NAM as an HDT. Since its discovery as one of the first effective TB therapies in the 1940s, it has been hypothesized that NAM exerts a host-directed effect^44,45,50^. We hypothesize that NAM exerts a host-directed effect as a precursor to NAD^+^. The effect of NAM is immediately dependent on enhanced glycolytic flux, as evidenced by inhibition of its effect by 2DG. It is worth noting that such a mechanism appears at odds with the proposed use of activators of the family of NAD^+^-consuming enzymes, sirtuins, as HDTs^51–53^. The host-directed effect of increasing the activity of these enzymes, however, is thought to be related to their activity as epigenetic regulators rather than their role as consumers of NAD^+^. Thus, while dramatically high concentrations of NAM were recently reported to have detrimental effects in cell culture, arguably owing to nonspecific inhibition of SIRT7^53^, NAD^+^ repletion through NAM supplementation in infected cells may well support the protective effects of sirtuins.

Despite its relatively early discovery, NAM appears to have been abandoned as a therapy during the golden age of antibiotic discovery for TB following a brief report suggesting it antagonizes isoniazid when used in combination^50,54^. While this may be true, the landscape of TB has shifted dramatically in the last 60 years, with an increase in the incidence of TB to over 10 million new cases annually and the development of resistance to the frontline drugs that displaced NAM. Here, we provide further validation of the host-directed effect of NAM, a mechanism for its activity, and a modern-day demonstration of its efficacy as a treatment for TB using two treatment regimens *in vivo*. Logistically, it satisfies many of the criteria for an optimal novel TB treatment regimen set forth by the WHO, given that it is inexpensive, orally bioavailable, shelf stable, and remarkably safe and tolerable. Additionally, the host-directed mechanism would make the development of bacterial resistance more difficult when compared to traditional antibiotics^55^. Finally, it is well studied and routinely used in humans for various indications^50,56–62^. Ultimately, these characteristics make NAM appealing as an old tool in new settings, including individual or community-level prophylaxis in high-incidence populations and for incorporation into drug regimens for isoniazid-resistant strains of *Mtb*.

Limitations of our study include that we primarily considered *Ldha* deletion as a specific approach to the NAD(H)-mediated manipulation of glycolysis. However, when pyruvate is a substrate, lactate is the product, and lactate has a variety of immunomodulatory effects in and beyond TB^30,63,64^. Given that we used a myeloid-specific knockout mouse, and lymphocytes make up the vast majority of cells within the lungs of both WT and *Ldha*^*LysM−/−*^ mice at all time points surveyed, we consider it unlikely that lactate played a greater role than the glycolytic defect we directly observed in myeloid cells.

In summary, we demonstrate the importance of two complimentary pathways responsible for the maintenance of NAD(H) homeostasis, lactate metabolism and NAD^+^ salvage, for host protection mediated by myeloid cells in TB. We observe that *Mtb*-infected macrophages exhibit an increased reliance on NAD^+^ salvage, which we expand to connect independent observations that *Mtb* infection diminishes NAD^+^ levels in macrophages^40,41^, *Mtb* decreases glycolytic capacity in infected macrophages^16,19,20^, and the NAD^+^ precursor NAM exerts a host-directed antimycobacterial effect^42–45^. Finally, we validate the antimycobacterial effect of NAM in an animal model of TB, providing rationale for its use as a new approach to TB prophylaxis or treatment.

## Acknowledgements

The authors wish to thank Drs. Pankaj Seth and Barbara Wegiel (Harvard Medical School) for providing *Ldha*^*LysM−/−*^ mice. This work was supported by the staff, management, and resources of the UAB Bioanalytical and Redox Biology Core, Heflin Genomics Core, Comprehensive Flow Cytometry Core, and the Southeastern Biosafety Laboratory Alabama Birmingham (SEBLAB), a NIAID-supported (UC6 AI058599) Regional Biocontainment Laboratory.

## Materials and Methods

### Human Subjects

The study of human lung pathology was approved by the University of KwaZulu-Natal Biomedical Research Ethics Committee (BREC, Class approval study number BCA 535/16). Patients undergoing lung resection for TB, their study protocol, associated informed consent documents, and data collection tools were approved by the UKZN BREC (Study ID: BE 019/13). Written informed consent was obtained from patients recruited from King DinuZulu Hospital Complex, a tertiary center for TB patients in Durban, South Africa.

### Animals

*LysM*^+/+^*Ldha*^fl/fl^ (*Ldha*^fl/fl^) and *LysM*^+/cre^*Ldha*^fl/fl^ (*Ldha*^*LysM*−/−^) mice on a C57BL/6 background were originally sourced from an independent investigator^30^ and maintained in pathogen-free facilities. C57BL/6J mice were obtained from the Jackson Laboratory. All studies used sex-matched, age-matched, littermates of both sexes. Littermates within each sex were cohoused until the start of experiments. For *in vivo* experiments, mice were aged 8-12 weeks old at the start, while, for the *ex vivo* generation of BMDMs, mice aged 8-16 weeks old were used. At the start of *in vivo* experiments, mice were housed under ABSL-3 conditions and monitored daily. All procedures and protocols were approved by the Institutional Animal Care and Use Committee of the University of Alabama at Birmingham.

### Bacteria

*Mtb* H37Rv was used in all experiments. Liquid cultures of *Mtb* were grown at 37 °C with shaking in Middlebrook 7H9 (Thermo Fisher Scientific, DF0713-17-9) broth supplemented with 0.02 % tyloxapol and albumin (Millipore Sigma, 3116956001), Dextrose (Thermo Fisher Scientific, DF0155-17-4), and saline (ADS) while solid cultures were grown at 37 °C standing on Middlebrook 7H11 agar (Thermo Fisher Scientific, L12203) supplemented with ADS and 0.2 % glycerol. Stocks of *Mtb* were generated from mid-log phase liquid cultures by 1:1 dilution with 50% glycerol in water and stored at −80°C for up to 6 months.

### Immunohistochemistry

Human lung samples were aseptically removed and fixed in 10% neutral buffered formalin (10% NBF). These samples were processed in a vacuum filtration tissue processor using a xylene-free protocol. Tissue sections were embedded in paraffin wax. Human lung tissue was cut into 2 μm-thick sections, mounted on charged slides, and heated at 56 °C for 15 minutes on a hotplate. Mounted sections were dewaxed in 2 changes of xylene followed by rinsing in 2 changes of 100% ethanol and 1 change of SVR (95%). Slides were then rinsed in tap water for 2 minutes followed by antigen retrieval via Heat Induced Epitope Retrieval (HIER) in trisodium citrate (pH = 6.0) for 30 minutes. Slides were cooled for 15 min and rinsed in tap water for 2 minutes. Endogenous peroxide activity was blocked by exposing tissue sections to 3% hydrogen peroxide (Novolink) for 10 min at room temperature (RT). Slides were then rinsed in PBST and blocked with protein block (Novolink) for 5 minutes at RT. Sections were incubated with primary antibody for lactate dehydrogenase (LDHA ab125683, abcam,1:500) followed by rinsing in PBST and incubated with the appropriate polymer (Novolink) for 30min at RT. Slides were then rinsed and stained with Diaminobenzidine (DAB) for 5 minutes, rinsed under running water, and counterstained with hematoxylin for 2 minutes. Slides were rinsed in tap water, blued in 3% ammoniated water for 30 seconds, rinsed in tap water, dehydrated in ascending grades of alcohol, cleared in xylene, and mounted with DPX (Distyrene, Plasticizer, and Xylene). Slides were scanned using a Hamamatsu NDP slide scanner (Hamamatsu NanoZoomer RS2, Model C10730-12) and its viewing platform (NDP.View2).

### Ex vivo experiments

For the generation of BMDMs, mice were euthanized according to institutional protocols. Femurs and tibias were isolated, and the bone marrow was flushed out through a cell strainer (70 μM pore size) with PBS supplemented with 5% FBS. Red blood cells were lysed with ACK lysis buffer, and the residual bone marrow cells were plated in tissue-culture-treated microplates at a density of 2 × 10^5^ cells per well in RPMI 1640 containing 2 mM glutamine (Thermo Fisher Scientific, 21875034), 10% FBS (Thermo Fisher Scientific, 10082147), 10 mM HEPES, and 5 ng/mL M-CSF (BioLegend, 576404; complete RPMI), as well as penicillin (100 units / mL) and streptomycin (100 units / mL), (AB/AM). Cells were differentiated in complete media with AB/AM and supplemented to 20 ng/mL M-CSF for 6 days. Media was changed once during this time, on day 3.

After 6 days, BMDMs were exposed to their indicated treatment condition in complete RPMI overnight (∼18 hours): IFNγ stimulation (10 ng/mL; BioLegend, 575304); supplementation with pyruvate or nicotinamide (NAM) (1 mM; Millipore Sigma); treatment with the LDHA/LDHB inhibitor, GSK 2837808A (5 μM; Tocris, 5189) or the NAMPT inhibitor FK866 (200 μM; Millipore Sigma, F8557); and/or *Mtb* infection at a multiplicity of infection (MOI) of 5:1. For *Mtb* infections, *Mtb* was quantified by optical density at a wavelength of 600 nm (OD_600_), assuming an OD_600_ of 1 corresponds to ∼10^8^ CFU. The required amount of liquid culture in the mid-log phase of growth or glycerol stock was passaged 5 times through a 27-gauge needle and added to complete RPMI (infection media). BMDMs were incubated with infection media at 37 °C and 5% CO_2_ for 4 hours, after which BMDMs were washed 3 times in PBS and returned to complete RPMI containing any other treatment conditions indicated within the experiment.

### Quantification of LDHA protein expression

Protein lysates from BMDMs were quantified by BCA analysis (Thermo Fisher Scientific, 23225), diluted to a concentration of 1 mg/mL, and mixed 1:1 with 2x Laemmli Buffer (Bio-Rad Laboratories, 161-0737). 10 μg of protein was loaded from each sample into separate wells of a 4-15% polyacrylamide gel (Bio-Rad Laboratories, 456-8084). Electrophoresis was performed in 1x Tris/Glycine Buffer (Bio-Rad Laboratories, 1610734) at 100 V, and protein was transferred to a PVDF membrane (Bio-Rad Laboratories, 1620177) overnight at 35V and 4 °C. Following the transfer, non-specific protein binding was blocked by incubation of the membrane in 5% non-fat milk for 30 minutes. The membrane was then incubated with the primary anti-LDHA antibody (ProteinTech, 19987-1-AP) diluted 1:1000 in 5% non-fat milk at RT for one hour. The membrane was washed three times in TBST and incubated with the HRP-conjugated goat-anti-rabbit secondary antibody (Kindle Biosciences, R1004) diluted 1:1000 in 5% non-fat milk at RT for one hour. The membrane was then covered in KwikKwant HRP Substrate Solutions A + B (Kindle Biosciences, R1004) and images on the KwikKwant Imager (Kindle Biosciences, D1001). Images were quantified with the ImageJ gel analysis tool^65^.

### LDH functional assay

LDH enzyme activity was determined by measuring the conversion of NADH to NAD^+^ in the presence of saturating concentrations of NADH and pyruvate. 10 μg of protein from separate BMDM cell lysates was loaded into separate wells and incubated in a final volume of 200 μL of PBS containing 10 mM pyruvate and 300 μM NADH (Acros Organics, 271102500) in an ultraviolet-transparent 96 well microplate (Thermo Fisher Scientific, 8404). NADH depletion was measured by the absorbance at 340 nm. To determine the rate of reaction, we performed a series of kinetic measurements with the minimal read time for the plate in the Synergy H1 Hybrid Multi-Mode Reader and Gen5 software (BioTek).

### Morphometric analysis

The open-source software QuPath (https://qupath.github.io)^33^ was used for morphometric analysis of whole-slide images. Stain vectors were adjusted for each image for optimal segmentation. Automated cell segmentation was performed for all tissue sections within using the “cell detection” tool following optimization of the segmentation parameters across representative sections from each experimental group. To determine the local cellular density measurement for each cell, we employed a modified version of the “create counts map.groovy” script published by the developer of the QuPath software (https://gist.github.com/petebankhead). In brief, the images were arbitrarily divided into many smaller regions (pixels) of equal size, and the number of nuclei within each region was determined. Pixels containing the edge of the tissue section were excluded from analysis. To improve on the discrete nature of this approach, the count of each pixel was adjusted to the mean values of its neighbors by applying a gaussian blur filter. The resulting value for each pixel was then assigned to the nuclei within that region. Data were exported and analyzed in R statistical software (see “Statistical analysis” section).

### *Aerosol infection with* Mtb

Mice within each experiment were infected as a cohort with *Mtb* H37Rv using an aerosol inhalation exposure system (Glas-Col) within ABSL-3 containment. *Mtb* was obtained from mid-log phase liquid cultures, washed once in PBS via centrifugation, and resuspended in 6 mL of PBS at an OD_600_ of 0.07. This suspension was then transferred to the nebulizer of the exposure system, allowing for exposure of the mice to aerosolized *Mtb*. 24 hpi, 3-4 mice were euthanized and the initial bacillary burden was determined as described in the section “Quantitation of bacterial burden.”

### Quantitation of bacterial burden

At indicated timepoints, mice were euthanized according to institutional protocols. Lungs and spleens were dissected and homogenized in 2 mL of PBS and serially diluted. 100 μL of each of dilution was plated on 7H11 agar as described in the section “Bacteria.” Colonies were counted after 21 days, and the total CFU per organ was determined by normalizing counts to dilution factor, volume plated, volume of the homogenate, and the proportion of the total organ plated.

### Flow cytometry

For flow cytometric analysis, the following anti-mouse antibodies and stain were acquired from BioLegend: PerCP-Cy5.5 anti-Ly6C (128012), APC-Cy7 anti-CD11b (101226), AF700 anti-Ly6G (127622), PE-Cy7 anti-F4/80 (123114), FITC anti-Siglec F (155504), BV785 anti-CD11c (117336), BV605 anti-CD45 (103115), BV421 anti-CD64 (107641), BV650 anti-IA/IE (107641), PE anti-CD24 (138503), and Zombie Aqua fixable viability dye (423101). Mice were euthanized at indicated timepoints according to institutional protocols. Immediately following sacrifice and thoracotomy, the pulmonary vasculature was perfused with PBS via the right ventricle. Lobes taken for cytometric analysis were minced in Dulbecco’s Modified Eagle Medium and incubated with Liberase (Roche) 2 mg/mL at 37 °C for 30 minutes. The resulting cell suspension was passed 5 times through a 20-gauge needle, 5 times through a 23-gauge needle and filtered through a cell-strainer with a 40 μm pore size. Cells were washed once in PEB buffer and stained with fixable live/dead stain for 30 minutes on ice. Cells were washed in PEB and stained for the indicated surface markers by staining cells with fluorophore-conjugated antibodies for 30 minutes at 4 °C. Cells were then fixed in 4% paraformaldehyde for 30 minutes, resuspended in PBS, and stored at 4 °C in the dark until they could be analyzed the following day. Flow cytometry acquisition was performed using the Attune NxT (Thermo Fisher Scientific), and analysis was performed with FlowJo software v10 (Tree Star).

### RNA sequencing and analysis

RNA was isolated from the lungs of *Mtb*-infected mice stored in RNAlater (Thermo Fisher Scientific, AM7024) at −80 °C using the RNeasy plus Mini Kit (Qiagen, 74134) according to the manufacturers protocol. Briefly, lung tissue was homogenized in RLT lysis buffer and passed through a genomic DNA eliminator column. The eluate was mixed with an equal volume of 70% ethanol and passed through an RNA-binding column. The column was washed once with RW1 buffer and twice with RPE, following which the membrane was thoroughly dried and the RNA was eluted in nuclease-free water.

mRNA-sequencing was then performed on the Illumina NextSeq500 as described by the manufacturer (Illumina Inc). RNA quality was assessed using the Agilent 2100 Bioanalyzer. RNA with a RNA Integrity Number (RIN) of ≥7.0 was used for sequencing library preparation. RNA passing quality control was converted to a sequencing ready library using the NEBNext Ultra II Directional RNA library kit with polyA selection as per the manufacturer’s instructions (New England Biolabs). The cDNA libraries were quantitated using qPCR in a Roche LightCycler 480 with the Kapa Biosystems kit for Illumina library quantitation (Kapa Biosystems) prior to cluster generation.

STAR (version 2.7.3a) was used to align the raw RNA-Seq fastq reads to the GRCm38 p6 Release M24 reference genome from Gencode^66^. Following alignment, HTSeq-count (version 0.13.5) was used to count the number of reads mapping to each gene^67^. Normalization and differential expression was then applied to the count files using DESeq2^68^. Further analysis of differentially expressed genes was performed as described in the section “Statistics and analysis.”

### Multiplexed cytokine analysis

Protein was isolated from the lungs of *Mtb*-infected mice stored in RNAlater at −80 °C by homogenization in RIPA buffer followed by 3 freeze-thaw cycles. Lysates were then centrifuged to remove debris, and the resulting supernatant was passed through a 0.22 μm spin column to removed debris and *Mtb*. The quantity of total protein in each sample was determined with the Pierce BCA Protein Assay Kit and each sample was diluted to 500 μg/mL.

Cytokine quantitation was performed with the Bio-Plex Pro Mouse Cytokine 23-plex Assay (Bio-Rad, M60009RDPD) according to the manufacturer’s instructions. In brief, samples were incubated with magnetic-bead-bound capture antibodies on a shaking platform at 850 RPM for 30 minutes followed by two washes in Bio-Plex wash buffer with a hand-held wash station. The process was repeated with biotinylated detection antibodies. Finally, PE-conjugated streptavidin was added to samples for a 10-minute shaking incubation. Samples were washed three times in Bio-Plex wash buffer, resuspended in assay buffer, and processed on the Magpix instrument platform with xPONENT software (Luminex).

### Extracellular flux analysis

Extracellular flux analysis was performed using the XFe96 Extracellular Flux Analyzer (Agilent), XFe96 Flux Packs (Agilent, 102416-100), and effector reagents purchase from Millipore Sigma. BMDMs were generated and treated in XFe96 cell culture plates as described in the section “*ex vivo* experiments.” A combination of the Glycolysis Stress Test and Mito Stress Test was performed as previously described^38^. Cells are incubated in glucose-free RPMI in a non-CO_2_ incubator the hour preceding the run. Initial ECAR measurements are used to determine non-glycolytic acidification. The first injection of glucose (final concentration: 10 mM) is followed by measurements to determine basal respiration (OCR) and glycolytic acidification (ECAR). The second injection of oligomycin (final concentration: 1.5 μM) is followed by measurements to determine ATP production by oxidative phosphorylation (OCR) and glycolytic capacity (ECAR). The third injection of FCCP and pyruvate (final concentrations: 1.5 μM and 1 mM, respectively) is followed by measurements to determine maximal respiration (OCR) and the spare respiratory capacity (OCR). Finally, the fourth injection of rotenone and antimycin A (final concentrations: 2.5 μM and 1.25 μM, respectively) is used to determine the non-mitochondrial oxygen consumption (OCR).

Additionally, the final injection also contained Hoescht 33342 (final concentration: 2 μg/mL; Thermo Fisher Scientific, 62249) to stain cell nuclei. Cell counts in each well were then determined with whole-well fluorescent imaging using a Cytation 5 Cell Imaging Multi-Mode Reader and Gen5 software (BioTek). In brief, images were acquired with the DAPI filter cube, 4x objective, and laser autofocus; stitched using the linear blend method; and processed using a combination of background flattening, blurring, and optimization of cellular analysis parameters. In instances of significant cell clumping (*i*.*e*., bacterial infection), cell count was determined by comparing mean fluorescent intensity to wells with an even distribution of cells (*i*.*e*. uninfected). ECAR and OCR values within each well were then normalized per 10,000 cells.

Bioenergetic parameters were determined from the ECAR and OCR values obtained from the glycolysis and mitochondrial stress tests as described elsewhere^11,12,38^. Beginning with parameters from the glycolysis stress test: Non-glycolytic acidification was determined by the third ECAR value following the initiation of the experiment (the ECAR value immediately preceding the injection of glucose). Glycolytic acidification was determined by subtracting non-glycolytic acidification from the third ECAR value following the injection of glucose (the ECAR value immediately preceding the injection of oligomycin). Glycolytic capacity was determined by subtracting non-glycolytic acidification from the third ECAR value following the injection of oligomycin (the ECAR value immediately preceding the injection of FCCP/pyruvate). Glycolytic reserve was determined by subtracting the value determined for glycolytic acidification from the value determined for glycolytic capacity.

Non-mitochondrial respiration was determined by the third OCR value following the injection of Rotenone/Antimycin A (the final OCR value in the experiment). Basal respiration was determined by subtracting non-mitochondrial respiration from the third OCR value following the injection of glucose (the OCR value immediately preceding the injection of oligomycin). Proton leak was determined by subtracting non-mitochondrial respiration from the third OCR value following the injection of oligomycin (the OCR value immediately preceding the injection of FCCP/pyruvate). ATP production was determined by subtracting proton leak from basal respiration. Maximal respiration was determined by subtracting non-mitochondrial acidification from the third OCR value following the injection of FCCP/pyruvate (the OCR value immediately preceding the injection of rotenone/antimycin A). Spare respiratory capacity (SRC) was determined by subtracting basal respiration from maximal respiration.

### Targeted metabolomics

Bone marrow cells were seeded in a 6-well microplate at a density of 3 × 10^6^ cells per well and differentiated as described in the section “ex vivo *experiments*.” Following the indicated treatment condition, cells were washed once in warm, glucose-free RPMI 1640 and incubated at 37 °C and 5% CO_2_ for 2 hours in glucose-free RPMI (Thermo Fisher Scientific, 11879020) supplemented with 10 mM ^13^C_6_-glucose (Cambridge Isotope Laboratories, CLM-1396-2) and dialyzed FBS (Thermo Fisher Scientific, 26400044). Following this incubation, cells were quickly washed three times with warm PBS and lysed by the addition of ice-cold extraction buffer (50% methanol in water) spiked with 20 ng/mL deuterated succinate (Millipore Sigma, 293075-1G) followed by scraping. Three technical replicate wells were pooled for each sample. Samples were subjected to three freeze/thaw cycles and then filtered through a 0.22 μm spin column. Protein content was estimated by the BCA assay, and samples were dried under vacuum for subsequent transport on dry ice. For liquid chromatography tandem mass spectrometry (LC-MS/MS), dried samples were reconstituted in 150 μL of water and filtered through a 0.22 μm spin column. 50 μL from each sample was mixed 1:1 with acetonitrile spiked with deuterated alanine for quantitation of amino acids. Standards for all metabolites analyzed were purchased (Millipore Sigma) and used as positive controls. Relative quantities of each metabolite were normalized to the internal standard, which is deuterated succinate for central carbon metabolites and deuterated alanine for amino acids, and to the protein concentration for that sample.

### Glucose uptake assay

Glucose uptake rates were determined with the Glucose Uptake-Glo Assay (Promega, J1341) according to the manufacturer’s instructions. Following stimulation, BMDMs were washed in PBS and incubated with 1 mM 2DG for 10 minutes, allowing for 2DG uptake and phosphorylation to 2DG-6-phosphate. After this incubation, stop buffer was added to each well followed by neutralization buffer. Finally, the 2DG-6-phosphate detection reagent was added, and samples incubated for 30 minutes at RT. Luminescence was quantified with an integration time of 1s using the Cytation 5 Cell Imaging Multi-Mode Reader (BioTek).

### NAD^+^/NADH quantification

NAD^+^ and NADH were quantified using the NAD/NADH-Glo Assay (Promega, G9071) according to the manufacturer’s instructions. BMDMs were washed once with PBS and media was replaced with 50 μL of PBS. 50 μL of 0.2 N NaOH with 1% CTAB was added to each well, and each sample was split into two 50 μL aliquots. To one group of aliquots, 25 μL of 0.4 N HCl was added, and both sets of samples were incubated for 15 minutes at 60 °C and 10 minutes at RT. 25 μL of 0.5 M Tris base was added to the HCL treated samples, while 50 μL of a 1:1 mixture of the HCl and tris solutions was added to untreated samples. 100 μL of NAD+/NADH-Glo detection reagent was then added to each well, and samples were incubated for 30 minutes at RT. Following incubation, luminescence was quantified with an integration time of 1 s using the Cytation 5 Cell Imaging Multi-Mode Reader (BioTek).

### Administration of compounds to mice

Nicotinamide (NAM) was administered to mice seven days per week via the drinking water at a concentration of 0.8% w/v, approximating a dose of 1 g/kg body weight/day. Therapeutic approaches consisted of a post-exposure prophylaxis model in which treatment was started at 3 days post infection and continued until sacrifice at 4 wpi, and a model for treatment of established infection in which treatment was started at 4 wpi and continued until 8 wpi.

### Statistical analysis

All statistical analyses were performed with R statistical software (version 4.0.2)^69^ in the RStudio integrated development environment (version 1.3)^70^. Unless otherwise specified, data were organized and graphed with the “tidyverse” collection of packages^71^. Heatmaps and hierarchical clustering was performed using the “pheatmap” package^72^. Gene set enrichment analysis was performed with the “fgsea” package to test for the enrichment of each of the 50 hallmark gene sets from the MSigDB, which collectively identify many of the most well-defined biological processes, among our differentially expressed genes ranked by fold-change^73^.

---

**Figure S1.**
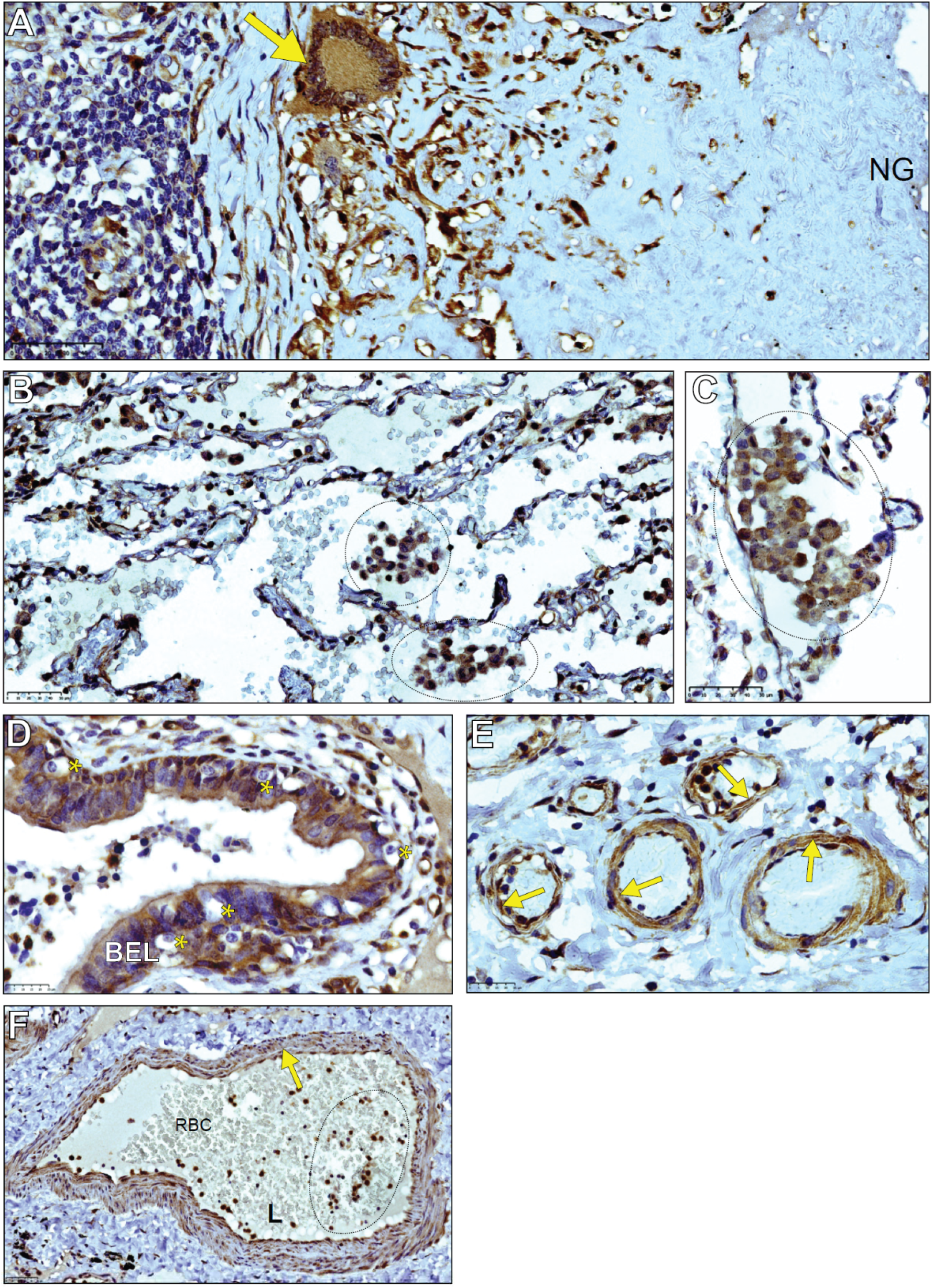
LDHA expression in alveolitis, giant cells, bronchial epithelial cells, and vasculature in the human TB lung. Low-**(A)** and **(B)** medium-power magnification images of LDHA immunostaining in leukocytes in the alveoli of a human TB patient. **(C)** High-power image of a giant cell stained for LDHA (yellow arrow) in the context of a necrotizing granuloma (NG). **(D)** shows the BEL stained for LDHA, as well as unstained cells (yellow asterisks) that represent mucus-producing cells. **(E, F)** Small **(E)** and large **(F)** vessel walls stained for LDHA expression (yellow arrows). **(F)** also shows robust LDHA staining in leukocytes within the lumen of the depicted vessel (dashed region).

**Figure S2.**
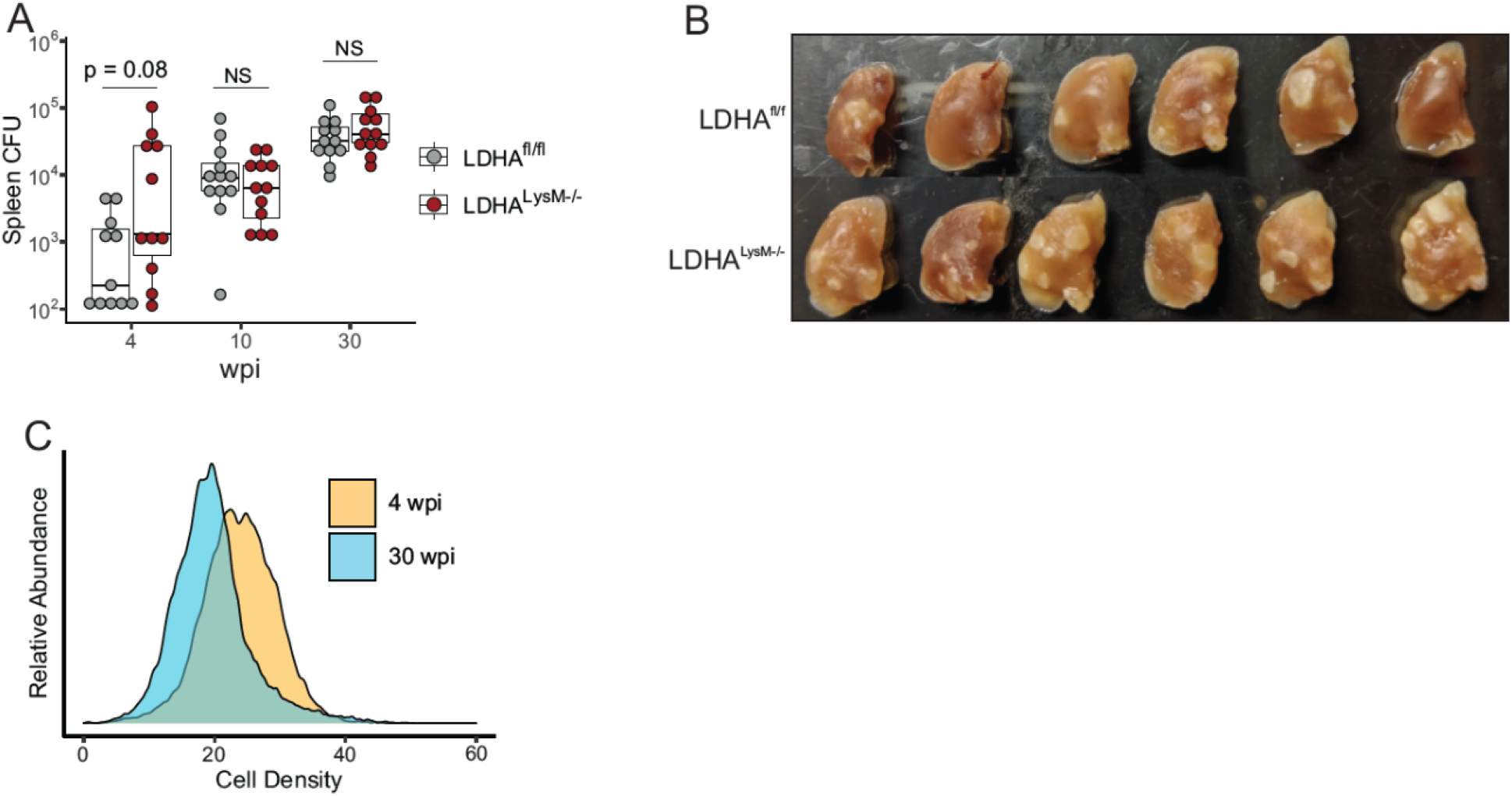
*Ldha*^*LysM−/−*^ mice are more susceptible to *Mtb* infection. **(A)** Box plot representing the burden of *Mtb* in the spleens of mice. Bottom, middle, and top horizontal lines for each condition represent the 25^th^, 50^th^, and 75^th^ percentile, respectively, while the whiskers extend from the edge of the box to the most distant point no further than 1.5 times interquartile range. Symbols represent biological replicates pooled from two independent experiments (n = 10-12 / group). **(B)** Gross pathology of formalin-fixed lungs from mice sacrificed at 30 wpi. **(C)** Cumulative histograms of all the cells within tissue sections of *Ldha*^fl/fl^ mice at 4 and 30 wpi, respectively. Statistical significance in (A) was determined by two-sample Wilcoxon rank-sum test.

**Figure S3.**
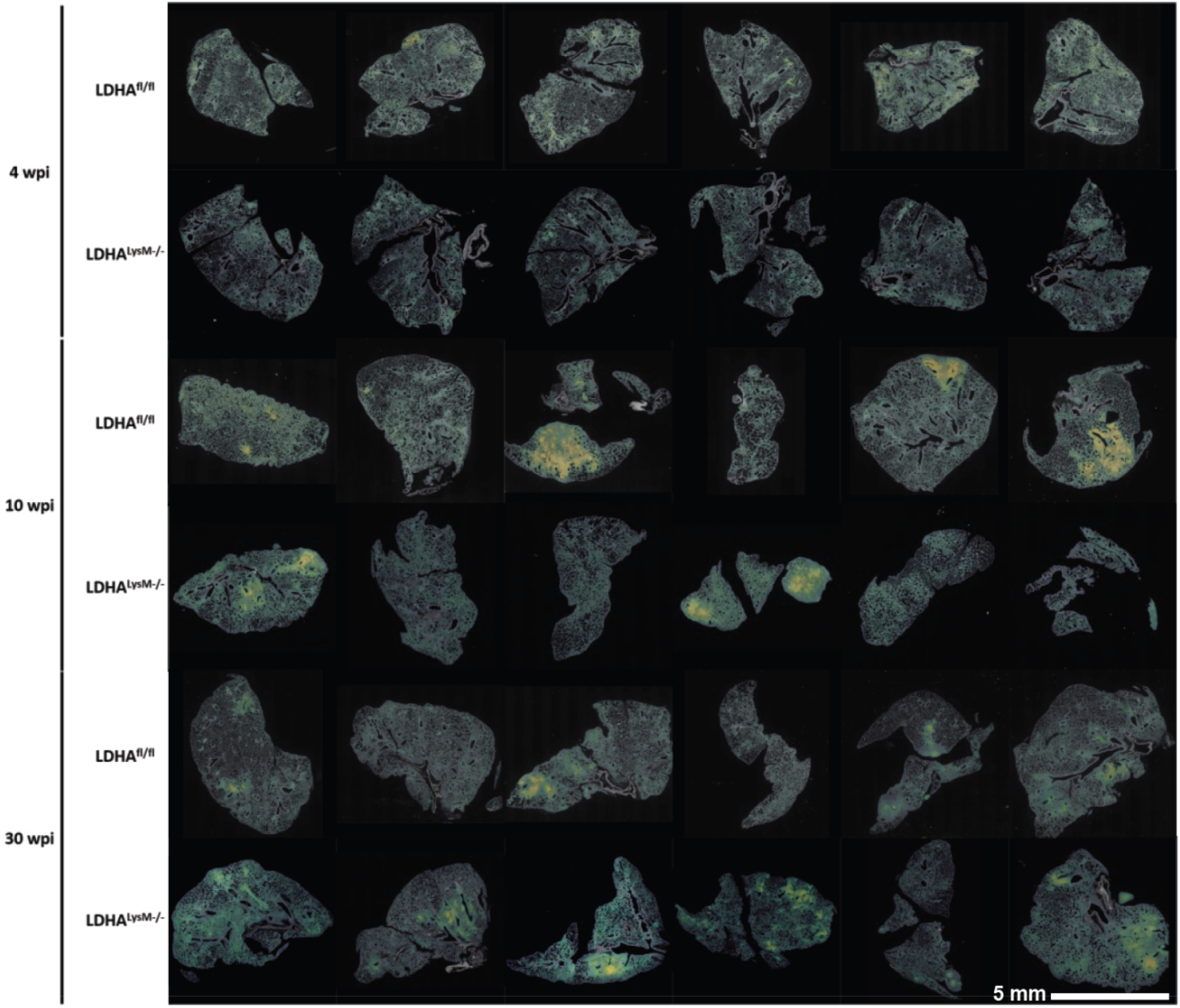
*Ldha*^*LysM*−/−^ mice exhibit slow-developing chronic inflammation following infection with *Mtb*. **(A)** Gray-scale rendering of the optical density of whole-tissue sections stained with hematoxylin and eosin. Nuclei were segmented based on optical density and pseudocolored with the viridis color scale based on local cell density, ranging from low (dark blue) to medium (green) to high (yellow). Scale bar (white) represents 5 mm.

**Figure S4.**
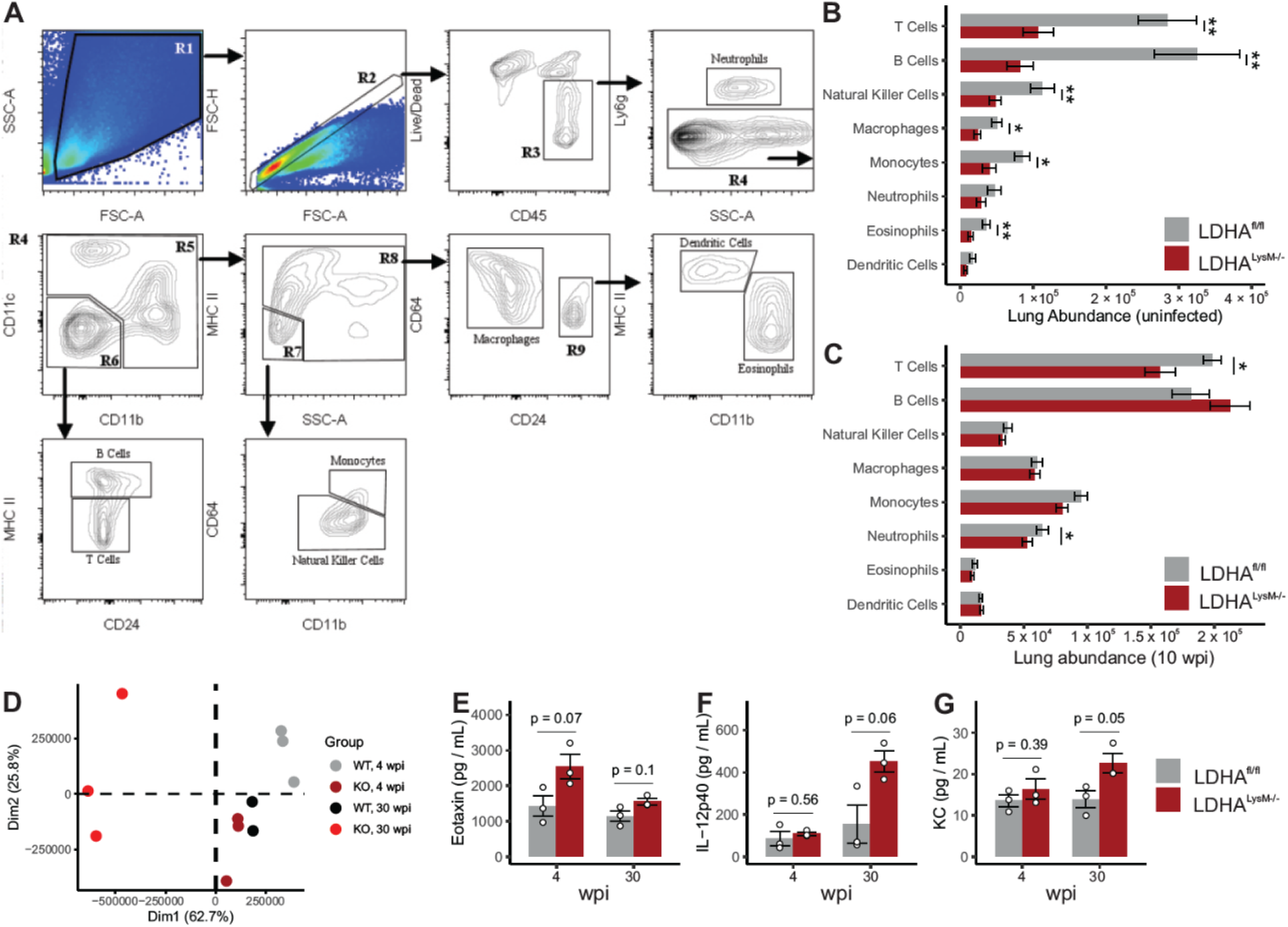
*Ldha*^*LysM*−/−^ mice exhibit dysregulated immunity during *Mtb* infection. **(A)** Gating strategy for multi-parameter flow cytometry. **(B, C)** Bar graph with error bars depicting the mean ± SEM for the absolute count of the indicated immune cell population in the lungs of *Mtb*-infected mice either uninfected **(B)** or at 10 wpi (C; n = 6/group). **(D)** Principal component analysis of the global transcriptome in the lungs of *Mtb*-infected mice. Symbols represent biological replicates. (**E-G)** Column graphs, error bars, and points representing the mean, SEM, and individual protein expression of biological replicates for the indicated cytokines (n = 3/group). Statistical significance was determined by the two-sample Wilcoxon rank-sum test **(B, C)** or two-sample t-test not assuming equal variance **(E-G)**. * p < 0.05, ** p < 0.01.

**Figure S5.**
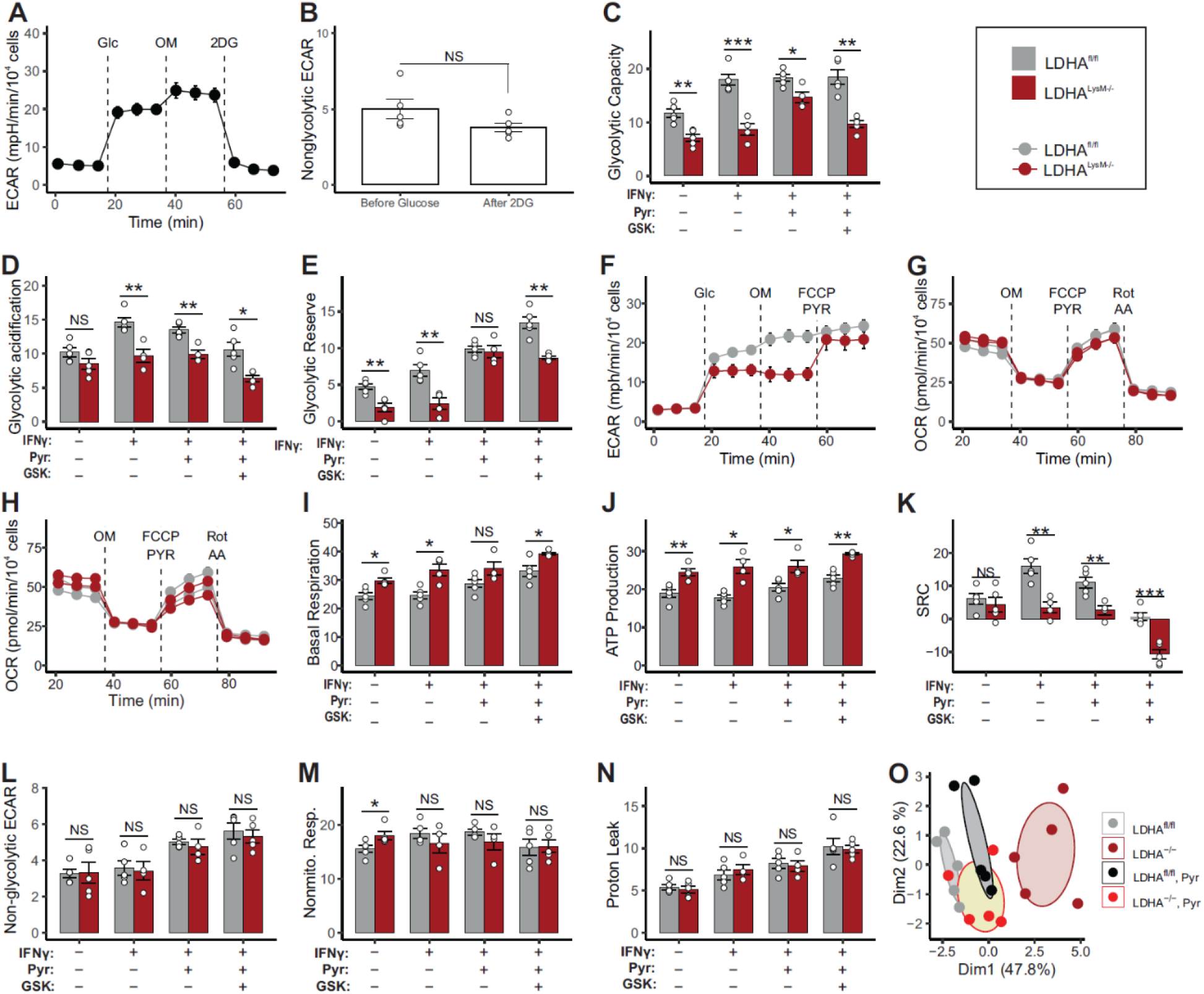
LDHA^−/−^ BMDMs stimulated with IFNγ exhibit diminished glycolysis and increased reliance on OXPHOS. **(A)** Line graph of the ECAR of BMDMs treated with 10 ng/mL IFNγ. Dashed lines in each panel represent the following injections in order: glucose, oligomycin, and 2DG. Symbols and error bars represent mean ±SEM of 5 biological replicates. **(B)** Columns, error bars, and symbols representing the mean, SEM, and individual values of biological replicates for nonglycolytic acidification as determined by the 3^rd^ read and 12^th^ read of panel A (n = 5/group). **(C-E)** Columns, error bars, and symbols representing the mean, SEM, and individual values of biological replicates for glycolytic capacity, basal glycolysis, and glycolytic reserve of BMDMs treated as indicated (n = 5/group). **(F)** Line graph of the ECAR of BMDMs treated with 10 ng / mL IFNγ. Dashed lines in each panel represent the following injections in order: glucose, oligomycin, and FCCP/pyruvate. Symbols and error bars represent mean ± SEM of 5 biological replicates. **(G, H)** Line graph of the OCR of BMDMs treated with D) 10 ng/mL IFNγ and 1 mM pyruvate (solid) or IFNγ alone (light) and E) IFNγ, pyruvate, and GSK 2837808A (solid) or IFNγ alone (light). Dashed lines in each panel represent the following injections in order: oligomycin, FCCP/Pyruvate, and Rotenone/Antimycin A. Points and error bars represent mean ± SEM of 5 biological replicates. **(I-N)** Columns, error bars, and symbols representing the mean, SEM, and individual values of biological replicates for basal respiration, ATP production by respiration, spare respiratory capacity, nonglycolytic acidification, nonmitochondrial respiration, and proton leak of BMDMs treated as indicated (n = 5/group). **(O)** Principal component analysis of the abundance of glycolytic and PPP intermediates in BMDMs stimulated with the indicated conditions. Symbols represent biological replicates and ellipses represent 95 % confidence intervals for each group. Statistical significance was determined by two-sample t-test without assuming equal variance **(B-E, I-N)**. * p < 0.05, ** p < 0.01, *** p < 0.001.

**Figure S6.**
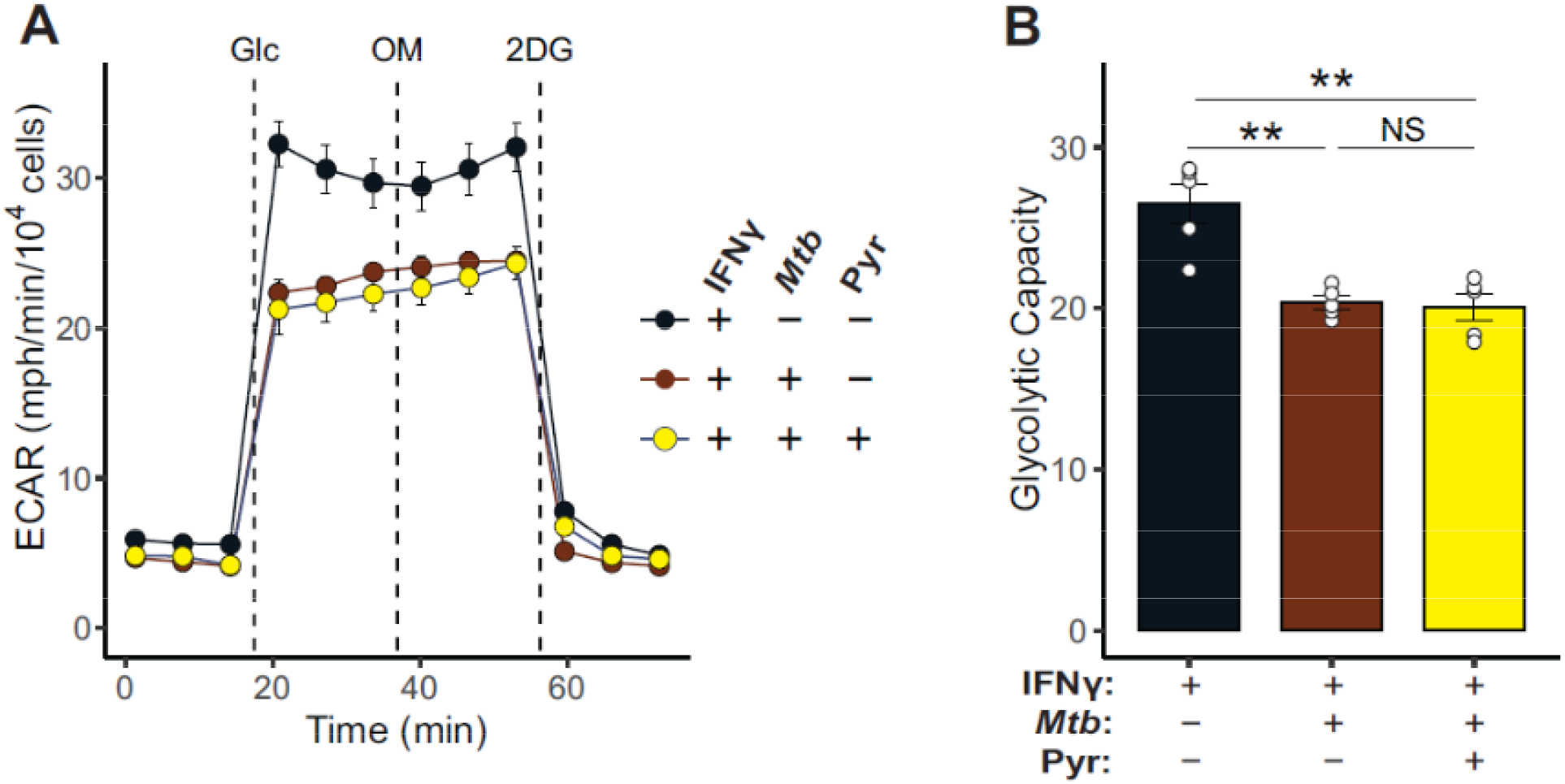
Pyruvate does not rescue *Mtb*-induced glycolytic defect in infected macrophages. (**A**) ECAR profiles for BMDMs exposed to combinations of IFNγ (10 ng/mL), *Mtb* (MOI 5:1), and pyruvate (1 mM) for 18 hours. Dashed lines represent injections of glucose (Glc), oligomycin (OM), and 2DG, respectively. Symbols and error bars represent mean ± SEM of 5 biological replicates. **(B)** Columns, error bars, and symbols representing the mean, SEM, and individual values for the glycolytic capacity determined from the profiles in panel A (n = 5/group).

